# Diversity and impact of single-stranded RNA viruses in Czech *Heterobasidion* populations

**DOI:** 10.1101/2024.05.01.591139

**Authors:** László Benedek Dálya, Martin Černý, Marcos de la Peña, Anna Poimala, Eeva J. Vainio, Jarkko Hantula, Leticia Botella

## Abstract

*Heterobasidion annosum* sensu lato comprises some of the most devastating conifer pathogens conifers. Exploring virocontrol as a potential strategy to mitigate economic losses caused by these fungi holds promise for the future. In this study, we conducted a comprehensive screening for viruses in a 98 *H. annosum* s.l. specimens from different regions of Czechia aiming to identify viruses inducing hypovirulence. Initial examination for dsRNA presence was followed by RNA-Seq analyses using pooled RNA libraries constructed from *H. annosum* and *Heterobasidion parviporum*, with diverse bioinformatic pipelines employed for virus discovery. Our study uncovered 25 distinct ssRNA viruses, including two ourmia-like viruses, one mitovirus, one fusarivirus, one tobamo-like virus, one cogu-like virus, one bisegmented narna-like virus and one segment of another narna-like virus, and 17 ambi-like viruses, for which hairpin and hammerhead ribozymes were detected. Coinfections of up to 10 viruses were observed in six *Heterobasidion* isolates, while another six harbored a single virus. 73% of the isolates analyzed by RNA-Seq were virus-free. These findings show that the virome of *Heterobasidion* populations in Czechia is highly diverse and differs from that in the boreal region. We further investigated the host effects of certain identified viruses through comparisons of the mycelial growth rate and proteomic analyses and found that certain tested viruses caused growth reductions of up to 22% and significant alterations in the host proteome profile. Their intraspecific transmission rates ranged from 0% to 33%. Further studies are needed to fully understand the biocontrol potential of these viruses *in planta*.

**Importance:**

- First report of a fusarivirus, a tobamo- and a cogu-like virus in *Heterobasidion*
- Certain viruses caused mycelial growth reduction in *H. annosum* host strains
- Viral infections lead to proteome changes in *Heterobasidion*

## 1. Introduction

The *Heterobasidion annosum* sensu lato species cluster comprises fungal root rot pathogens in the temperate and boreal regions of the Northern Hemisphere (Garbelotto and Gonthier 2013). These basidiomycetes represent a major threat to intensively managed forest stands, while playing a subordinate role in natural ecosystems (Shaw et al. 1994; Dálya and Sedlák 2020). Financial losses attributable to the pathogen in the European Union are approximately 1.4 billion € annually, inflation-adjusted from Woodward et al. (1998). Heterobasidion root rot has been a serious problem in planted Norway spruce (*Picea abies*) and Scots pine (*Pinus sylvestris*) stands in Czechia ever since forest management was introduced (Černý 1989; Sedlák and Tomšovský 2014). In 2020, the volume of salvage fellings in Czechia rose to an unprecedented 34 million m^3^, mostly due to *Ips typographus* as the main disease agent (Ministry of Agriculture of the Czech Republic 2021). Despite the extent of damages, no practical measures have been implemented in Czechia to control the spread of *H. annosum* s.l. because the problem generally receives little attention.

Preventive control, i.e. chemical (urea, borax) or biological (antagonistic fungus *Phlebiopsis gigantea*) stump treatments against aerial spore infection should be a high priority in disease management strategies against *Heterobasidion* spp. (Gonthier and Thor 2013). However, this is of little avail in many regions due to the nearly ubiquitous occurrence of the pathogen (Dálya et al. 2022). Currently available curative control methods are rotation of tree species at a site and stump removal (Garbelotto and Gonthier 2013). Change of the tree species is not always possible due to site characteristics or economic considerations. Stump harvesting removes only a fraction of the pathogen inoculum (Piri and Hamberg 2015) and may lead to several negative environmental impacts (Walmsley and Godbold 2010). The eradication of the fungus, once it has established itself in a stand, can be a very lengthy, laborious and costly process with varying success rates. During the past 25 years, the possibility of using mycoviruses as biocontrol agents was being explored in Sweden (e.g., Ihrmark et al., 2001) and Finland (e.g., Vainio et al., 2018) as a fundamentally new solution to limit *Heterobasidion* damages.

Viruses that inhabit fungi, i.e. mycoviruses, are hosted by a wide range of fungal taxa (Hough et al. 2023). The high phylogenetic diversity of mycoviruses is reflected by their current classification into 10 double-stranded (ds) RNA families, one unclassified dsRNA genus, 15 single-stranded (ss) RNA families and one ssDNA family (https://ictv.global/), and many novel taxa are awaiting description. The discovery rate of new viruses is exponentially accelerated by high-throughput sequencing approaches (Edgar et al. 2022). The incidence of mycoviral infections in plant-associated fungi ranges from a few % to over 90% (Hillman et al. 2018). Most of these virus/host interactions are cryptic, but beneficial or harmful effects are also known. Mycoviruses that induce hypovirulence in phytopathogenic fungi may be utilized as biological control agents. The best-known example is Cryphonectria hypovirus 1 (CHV1), which was successfully applied as a treatment against chestnut blight in Europe (Heiniger and Rigling 1994). In addition, virus-induced hypovirulence has been shown in *Rosellinia necatrix*, *Sclerotinia sclerotiorum*, *Botryosphaeria dothidea* and *Fusarium* spp., among other fungal pathogens (García-Pedrajas et al. 2019).

The *Heterobasidion* genus hosts a diverse mycovirus community with worldwide distribution. Viruses with dsRNA genomes infect approximately 15% of *H. annosum* s.l. isolates in Europe and Western Asia (Ihrmark 2001; Vainio et al. 2011a). The most common mycovirus in *Heterobasidion* spp., Heterobasidion RNA virus 6 (HetRV6; *Curvulaviridae* family), is responsible for more than 70% of all dsRNA infections (Vainio et al. 2012). HetRV6 was found in many different *Heterobasidion* species of distant origins (e.g., Europe, Siberia, USA) and showed a high degree of geographic and host-related differentiation (Vainio et al. 2012). Members of *Partitiviridae* are also widespread and highly diverse. So far, 21 different partitiviruses have been described in *Heterobasidion* spp. and all belong to the *Alphapartitivirus* or *Betapartitivirus* genera (Sutela et al. 2021). Mitoviruses (*Mitoviridae* family) were detected by small RNA deep sequencing and RNA-Seq in *Heterobasidion parviporum* and *H. annosum* (Vainio et al. 2015a; Vainio 2019). More recently, further positive-strand RNA viruses, representing the *Narnaviridae* and *Botourmiaviridae* families were discovered in *H. parviporum*, as well as members of proposed new phylum *Ambiviricota* (proposed families *Dumbiviridae* and *Trimbiviridae*) with ambisense RNA genomes (Sutela et al. 2021; Turina et al. 2023).

Ample evidence shows that virus transmission is possible within and between *Heterobasidion* species both *in vitro* (Ihrmark et al. 2002; Vainio et al. 2010; Vainio et al. 2011b; Vainio et al. 2011a; Vainio et al. 2012; Vainio et al. 2018; Kashif et al. 2019; Hantula et al. 2020) and in natural conditions (Vainio et al. 2013; Vainio et al. 2015b; Sutela et al. 2021; Piri et al. 2023). Pre-existing viruses in either the donor or the recipient mycelium may help or hinder the transmission of other viruses (Vainio et al. 2015b; Kashif et al. 2019; Hantula et al. 2020). While most viruses of *Heterobasidion* are considered more or less latent, both negative and slightly positive interactions with the fungal hosts have been reported (Ihrmark et al. 2004; Vainio et al. 2010; Vainio et al. 2012; Hyder et al. 2013; Jurvansuu et al. 2014; Vainio et al. 2018; Kashif et al. 2019). The same mycovirus can exhibit differential effects on its hosts depending on the particular host strain and environmental conditions (Vainio et al. 2010; Hyder et al. 2013). Laboratory and field experiments with Heterobasidion partitivirus 13 from *H. annosum* (HetPV13-an1) turned out promising for the development of a future biocontrol product (Vainio et al. 2018; Kashif et al. 2019; Hantula et al. 2020). By affecting the transcription of 683 genes (representing 6% of the host genome), the virus influenced the cell cycle, hampered the uptake of energy, caused metabolic changes, limited the growth of host isolates and altered mycelial morphology (Vainio et al. 2018). As a result, the virulence of its natural host, as well as that of the recipient *H. parviporum* strains reduced considerably. Very recently, HetPV13-an1 was shown to enhance the biocontrol effect of *P. gigantea* when applied to pine stumps infested by *H. annosum* (Piri et al. 2023).

As previous research focused on boreal forests, limited data is available about viral communities hosted by Central European *Heterobasidion* strains. However, of dsRNA viruses HetRV6 is known to occur in Austria and Poland (*Heterobasidion abietinum, H. parviporum* and *H. annosum*), and the partitiviruses HetPV11 (several isolates), HetPV12- an1, as well as the putative biocontrol virus HetPV13-an1 originate from Poland (*H. annosum*). We hypothesized that Czech *Heterobasidion* isolates harbor similar viruses. The specific aims of this study were to: a) collect ca. 100 strains of *H. annosum* s.l. from different regions of Czechia; b) screen the gathered isolates for virus presence by both traditional and state-of-the-art methods; c) describe the putative mycoviruses and characterize their genome; d) investigate the effects of selected viruses on the phenotype of their host. Depending on the outcomes of these experiments, the long-term objective is to explore the possibilities of virocontrol against Heterobasidion root rot with virus strains naturally distributed in Czechia. The present work might represent an early step towards restoring the health of the remaining conifer stands infested by *Heterobasidion*.

## 2. Materials and methods

### 2.1. Origin of the fungal isolates

Strains of *H. annosum* s.l. were collected by arbitrary sampling throughout Czechia, with an emphasis on managed stands of Norway spruce and Scots pine. Fruiting bodies were gathered from substrates (decayed stumps, logs, roots or the ground) located at least 30 m apart, to minimize the chance of repeated sampling of the same genotype (Stenlid 1985). Pure cultures were isolated from basidiocarps and grown on 2% malt extract agar (MEA; HiMedia, India) at 18°C. One genotype was obtained from a wood disc used as a spore trap, by isolating an emerging conidiophore. In total, 96 *Heterobasidion* strains were collected from Czechia, and two Slovakian strains were included in the study as well (Supplemental Table S1).

Isolates were identified to the species level as described earlier (Dálya et al. 2022). For isolates collected in 2018–19, the dilution protocol of the Phire™ Plant Direct PCR Kit (Thermo Fisher Scientific, Waltham, MA, USA) was followed with 2 µL supernatant as template. Mycelia were subcultured onto MEA covered by a thin cellophane membrane, incubated at 22°C in the dark for 3 weeks, harvested into sterile 50-mL Falcon tubes, lyophilized at −55°C for 20 h and pulverized with two autoclaved 10-mm steel balls by vortexing for 2 min.

### 2.2. RNA extractions

The presence of dsRNA elements in the isolates was examined using CF11 cellulose affinity chromatography (Morris and Dodds 1979), modified by Tuomivirta et al. (2002), using 2 g of fresh mycelium. The size of the dsRNAs was estimated by electrophoresis using a 1.2% agarose gel. Total RNA was extracted by the Quick-RNA™ Fungal/Bacterial Miniprep Kit (Zymo Research, Irvine, CA, USA) from the fungal cultures grown and prepared as described in the previous section. Homogenization was done by shaking for 2 min at 30 Hz in a Mixer Mill MM 400 (Retsch, Haan, Germany). The quality of total RNA was visually assessed using a 1.2% agarose gel. RNA concentrations were measured by a Qubit^®^ 2.0 Fluorometer (Invitrogen). The RNA solutions were stored at −80°C.

### 2.3. RNA Sequencing

Three pools consisting of total RNA of 16 *H. annosum* strains, another 16 *H. annosum* strains and 13 *H. parviporum* strains (Supplemental Table S1) were treated with the TURBO DNA-free™ Kit (Thermo Fisher Scientific) and sent to SEQme s.r.o. (Dobříš, Czechia). Ribosomal RNA was removed using the NEBNext^®^ rRNA Depletion Kit (Human/Mouse/Rat). The cDNA libraries were constructed using the NEBNext^®^ Ultra II Directional RNA Library Prep Kit for Illumina^®^ and sequenced in paired-end (2×150 bp) on a NovaSeq 6000 (Illumina). After the confirmation of multiple virus infections in *H. annosum* isolate 1987, RNA-Seq was repeated for this single strain as described above, with the following exceptions. Total RNA was isolated using RNAzol^®^ RT (Sigma-Aldrich, Steinheim, Germany) with homogenization lasting 1 min. DNase treatment was omitted and RNA-Seq was performed by the Institute of Applied Biotechnologies a.s. (Prague, Czechia) with a read length of 2×151 bp.

### 2.4. Bioinformatics

Two approaches were combined for *in silico* data mining for viruses in the NGS datasets, each of which consisted of ca. 600 million reads. The quality of raw reads was assessed using FastQC-0.11.8 (https://www.bioinformatics.babraham.ac.uk/projects/fastqc, accessed on 11 April 2022) and adapter sequences clipped with cutadapt-3.4 (Martin 2011). The first bioinformatic pipeline consisted of *de novo* assembly with Trinity-2.11.0 (Grabherr et al. 2011) and searching for similarities of the obtained contigs to custom virus and host protein and nucleotide reference databases obtained from NCBI, using the BLASTX and BLASTN algorithms of the BLAST+-2.10.0 program (Camacho et al. 2009). The E value threshold was 1e-5. Contigs showing homologies to viral proteins or genomic sequences were selected for further analysis, while contigs with homologies to the host genome were excluded.

In the second pipeline, reads mapped to the host genome with BWA-0.7.12-r1034 MEM algorithm (Li 2013) were removed using SAMtools-1.9 (Danecek et al. 2021). The BamTools-2.5.1 suite (Barnett et al. 2011) was used for generating mapping statistics and for file format conversion. Mapping quality was also evaluated by QualiMap-2.2.2, BAM QC mode (Okonechnikov et al. 2016). The host-unmapped reads were aligned to a nucleotide database of *Heterobasidion* viruses with BWA MEM. Alignments were visualized in Tablet-1.12.02.08 (Milne et al. 2013). Random sequencing errors in the host-unmapped reads were fixed with Rcorrector-1.0.4 (Song and Florea 2015). Read pairs where at least one read was deemed unfixable were discarded using the “FilterUncorrectabledPEfastq.py” python script (https://github.com/harvardinformatics/TranscriptomeAssemblyTools/blob/master/FilterUncorrectabledPEfastq.py, accessed on 11 April 2022). The *de novo* assembly of host-unmapped reads and the subsequent similarity searches were conducted as in the first pipeline.

The lists of virus candidate contigs identified via both pipelines, together with their best hits in BLASTX and BLASTN were examined further. Contigs representing potential mycoviruses, based on our subjective evaluation, were grouped by their best hit in BLASTX. Within each of these groups, one contig deemed the most representative of the putative mycovirus according to its length and alignment parameters was imported to Geneious Prime^®^ 2021.1.1 (https://www.geneious.com). All contigs from both Trinity assemblies of all three pooled datasets were mapped against each selected contig using Geneious mapper with medium sensitivity. After the manual correction of misalignments, the resulting consensus sequences were saved and used in further analyses.

The screening of small self-cleaving ribozymes was carried out using the INFERNAL software (Nawrocki and Eddy 2013), which identifies RNA elements using secondary structure and nucleotide covariation modeling. Covariance models for the small ribozymes recently described from diverse ambiviruses, mitoviruses and other viroid-like agents were used for the searches (see Forgia et al. (2023)).

### 2.5. RT-PCR screening

cDNA was synthesized from the total RNA samples using either the ProtoScript^®^ II First Strand cDNA Synthesis Kit (New England Biolabs, Ipswich, MA, USA), following the standard protocol with Random Primer Mix, or the High-Capacity cDNA Reverse Transcription Kit (Thermo Fisher Scientific). To reveal the identity of virus-hosting *Heterobasidion* strains, RT-PCRs of the strains belonging to the RNA pool in which the putative virus was detected (Supplemental Table S1) were performed with virus-specific consensus primers (Supplemental Table S2). These primers were designed by Primer3-2.3.7 under Geneious to amplify a fragment within an ORF of putative viruses. Amplifications were carried out by the Phire™ Green Hot Start II PCR Master Mix (Thermo Fisher Scientific) or the OneTaq^®^ Quick-Load^®^ 2X Master Mix with Standard Buffer (New England Biolabs) with 2.5 µL cDNA in 25 µL reaction volume. PCR products were electrophoresed in a 1% agarose gel (Supplemental Fig. S1). When multiple bands were visible, the targeted one was purified from a gel using the NucleoSpin^®^ Gel and PCR Clean-up (MACHEREY-NAGEL, Düren, Germany) or the Monarch^®^ DNA Gel Extraction Kit (New England Biolabs). In a few cases, a second PCR with 0.5–1 µL of the PCR product as template was necessary to obtain a band with sufficient concentration. Target amplicons were sequenced by Eurofins Genomics (Ebersberg, Germany).

DNA was extracted from lyophilized mycelium of isolate 1987 with the DNeasy^®^ Plant Mini Kit (QIAGEN, Valencia, CA, USA) and used as template in a PCR to rule out the integration of one movement protein (MP) encoding viral segment in the host genome (see section 3.8). Additionally, 85 *H. annosum* s.l. isolates (Supplemental Table S1) were screened for the presence of HetRV6 with specific primers designed by Vainio et al. (2012). Here, 2 µL cDNA was used as template in RT-PCR; the rest of the workflow was as described above.

### 2.6. Confirmation of the 3’ end of an incomplete viral genome

Rapid amplification of cDNA ends (RACE) of the incomplete RNA-dependent RNA polymerase (RdRp) encoding region at the 3’ terminus of a putative linearized ambi-like virus was done following the protocol of the SMARTer^®^ RACE 5’/3’ Kit (Takara Bio USA, Mountain View, CA, USA), employing a specific primer designed by Primer3-2.3.7 under Geneious (Supplemental Table S2). PolyA RNA was purified with the RNA Clean & Concentrator-5 Kit (Zymo Research) and quantified using a BioSpec-nano spectrophotometer (Shimadzu, Japan). The RACE product was cloned using the In-Fusion^®^ HD Cloning Kit (Takara Bio USA) following the producer’s instructions and modifications as described by Botella et al. (2020).

### 2.7. Genetic variability and phylogenetic analyses

For calculating the genetic distance between mycovirus strains, pairwise (pw) sequence alignments of Sanger sequenced amplicons, as well as their translations, were done by MAFFT-7.490 (Katoh and Standley 2013) under Geneious. Maximum likelihood phylogenetic trees based on the amino acid (aa) sequences of either the RdRp or the MP were constructed using the bootstrapping algorithm implemented in MEGA11 (Tamura et al. 2021). The tree for the *Virgaviridae* family was built based on the codon-aligned nucleotide sequences of the replication proteins and RdRps.

### 2.8. Virus elimination

To assess the effect of mycoviruses on their host, it was necessary to create isogenic lines of selected fungal isolates with and without virus(es). In an attempt to eliminate viruses, 10 monohyphal cultures of three naturally infected *H. annosum* isolates (1987, 1989, 2072) were generated as follows. Hyphal tips of isolates grown on 3% water agar were picked with a Pasteur pipette under a microscope and inoculated onto MEA plates. Subcultures of the resulting single hyphal isolates were cultivated on cellophane-covered MEA plates for ca. 2 weeks. Total RNA was extracted from the harvested mycelia by the E.Z.N.A.^®^ Fungal RNA Mini Kit (Omega Bio-tek, Norcross, GA, USA), following the standard protocol. The fungal tissue was disrupted by a FastPrep-24™ Classic homogenizer (MP Biomedicals, Santa Ana, CA, USA) and 1–2-mm quartz sand grains. The RNA concentration and integrity were measured with a NanoDrop™ One^C^ spectrophotometer (Thermo Fisher Scientific). cDNA was synthesized from 3 µg RNA using RevertAid™ Reverse Transcriptase (Thermo Fisher Scientific) and random hexamer primers with a denaturation step at 98°C for 5 min. The presence of viruses in the single hyphal cultures was confirmed via RT-PCR, using DreamTaq™ Green DNA Polymerase (Thermo Fisher Scientific) with 1 µL cDNA in 20 µL reaction volume. PCR products were electrophoresed in a 1% agarose gel. When multiple bands appeared, the targeted one was purified from a gel using the E.Z.N.A.^®^ Gel Extraction Kit (Omega Bio-tek). Target amplicons were sequenced at Macrogen Europe (Amsterdam, The Netherlands).

As the above method was only partially successful, thermal treatment was applied for virus removal (Vainio et al. 2010; Vainio et al. 2018). Four monohyphal cultures derived from isolate 1987, each having retained different combinations of viruses, and the original isolates 1987, 1989 and 2072 were cultivated on MEA at 32°C for 1 week and at 33°C for another week. Subcultures were taken at the end of both weeks, allowed to recover to a normal growth rate at 20°C and subjected to RT-PCR screening.

### 2.9. Horizontal transmission

Virus transmission was expected to occur via hyphal anastomosis in dual cultures. Each of three virus-infected (1987, 1989, 2072) *H. annosum* isolates was paired with three virus-free (1985, 2013, 2052) *H. annosum* strains on 9-cm MEA plates and incubated at 22°C in the dark for 1 month. Small agar pieces taken from the recipient side were transferred to new MEA plates partially covered by cellophane. The sampling spots were located: 1) within 1 cm from the interaction zone separating the donor and the recipient strain; 2) at ca. 2 cm from the interaction zone (Fig. 1A). After 2–5 weeks, fresh mycelium was ground in liquid N_2_ using sterile mortar and pestle and total RNA was extracted using RNAzol^®^ RT. cDNA synthesis and RT-PCR were conducted as for the initial screening. The whole experiment was repeated using a replicate plate of the same pairings incubated for 3 months. This time, the following spot was also sampled: 3) far from the interaction zone, near the edge of the Petri dish (Fig. 1B).

**Fig. 1.**
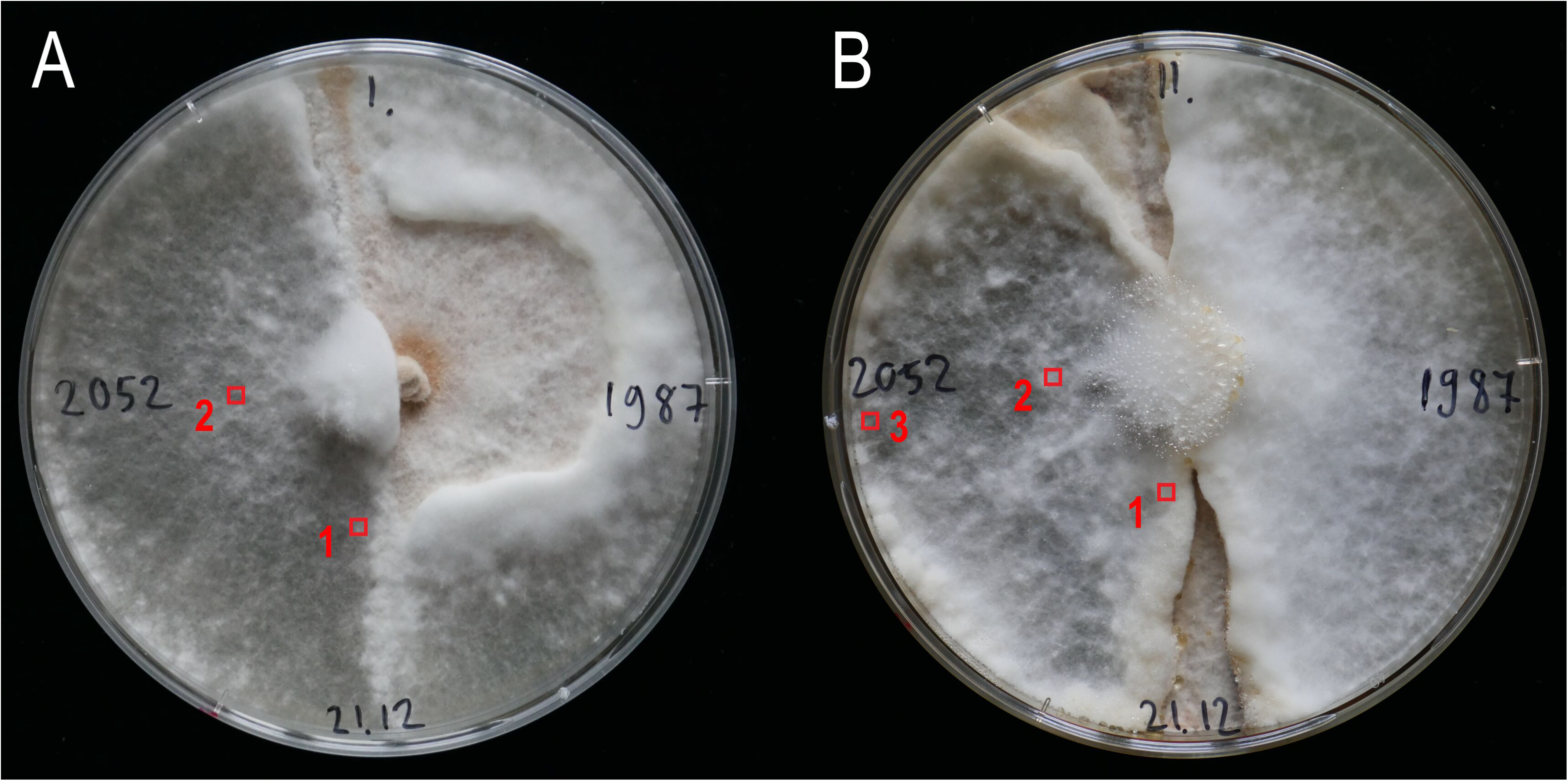
Arrangement of dual cultures used to transmit mycoviruses from donor (1987) into recipient (2052) *H. annosum* strains, showing the location of sampling spots 1 month (**A**) and 3 months (**B**) post-inoculation

The genetic identity of the virus-infected recipient isolates was verified with random amplified microsatellite (RAMS) fingerprinting with CCA primer (Hantula et al. 1996). DNA was isolated from fresh mycelium using the DNeasy^®^ Plant Mini Kit with 2×50 µL elution volume. Amplifications were performed in two repetitions by the OneTaq^®^ Quick-Load^®^ 2X Master Mix with Standard Buffer, with 2 µM primer and 0.5 µL DNA in 25 µL reaction volume. Thermal cycling was conducted as follows: 4 min at 94°C, followed by 35 cycles of 30 s at 94°C, 45 s at 55°C and 2 min at 68°C, with a final extension of 5 min at 68°C. PCR products were electrophoresed in a 1.8% agarose gel.

### 2.10. Growth rate measurements

Fifteen *H. annosum* isolates were selected for growth rate comparison *in vitro*. Circular agar plugs with fresh mycelium were inoculated onto the center of 9-cm MEA plates (five replicates) and incubated at 22°C in the dark. The mycelial edge was marked at the bottom of the plates 3 and 5 days post-inoculation (dpi) and subsequently photographed. The area of fungal colonies was measured using ImageJ-1.53v (Schneider et al. 2012). Comparison of the absolute growth of *a priori* determined pairs of isolates 3 and 5 dpi was done by Student’s *t*-test. The levels of significance were defined as *p*<0.05 and *p*<0.001. Statistical analyses and data visualization were performed in RStudio (https://www.r-project.org).

### 2.11. Proteomic analysis

The cellular response of *H. annosum* to mycoviral infections was investigated by proteomics. Cellophane-covered MEA plates (four replicates) were inoculated with the same fungal strains as in the growth rate experiment. After incubation at 22°C in the dark for 12 days, mycelia were harvested into sterile 15-mL Falcon tubes and lyophilized at −55°C for 4 h. Total protein extracts were prepared as previously described (e.g., Dufková et al., 2023) and portions of samples corresponding to 5 µg of peptide were analyzed by nanoflow reverse-phase liquid chromatography-mass spectrometry using a 15 cm C18 Zorbax column (Agilent, Santa Clara, CA, USA), a Dionex Ultimate 3000 RSLC nano-UPLC system, and the Orbitrap Fusion Lumos Tribrid Mass Spectrometer equipped with a FAIMS Pro Interface. All samples were analyzed using FAIMS compensation voltages of −40, −50, and −75 V. The measured spectra were recalibrated, filtered (precursor mass—350–5000 Da; S/N threshold—1.5), and searched against the *Heterobasidion annosum* v2.0 (Olson et al. 2012) and common contaminants databases using Proteome Discoverer 2.5 (Thermo Fisher Scientific, algorithms SEQUEST; MS Amanda, Dorfer et al., 2014). The quantitative differences were determined by Minora, employing precursor ion quantification followed by normalization (total area) and calculation of relative peptide/protein abundances. The reported statistical tests were generated and implemented as follows using the default and recommended settings. The reliability of the protein identifications was assessed in Proteome Discoverer 2.5. The Student’s *t*-test was calculated using MS Excel. For the ANOVA with Tukey’s HSD and the Kruskal-Wallis tests, the Real Statistics Resource Pack software for MS Excel (Release 6.8; Copyright 2013–2020; Charles Zaiontz; https://www.real-statistics.com) and MetaboAnalyst 5.0 (Pang et al. 2021) were employed. OPLS and the VIP was performed in SIMCA 14.1 (Sartorius, Goettingen, Germany). Significant differences refer to *p*<0.05.

## 3. Results and discussion

### 3.1. Virus detection and discovery

The applied methodology allowed for the reliable detection of known viruses, as well as the discovery of novel viruses. No dsRNA elements were found in our collection of 98 *Heterobasidion* isolates analyzed using cellulose chromatography. In agreement with this result, HetRV6 was absent in a subset of 85 isolates screened with RT-PCR. Altogether, 25 ssRNA viruses were discovered in the RNA-Seq datasets comprising altogether 32 *H. annosum* and 13 *H. parviporum* isolates (Table 1). 27% of these isolates were infected by at least one virus. All detected viruses were hosted by *H. annosum*, except one strain of the recently described Heterobasidion ambi-like virus 3 (HetAlV3) (Sutela et al. 2021), which dwelt in *H. parviporum* (Table 1). The presence of each virus was verified in up to five host strains. Six isolates harbored a single virus, while coinfections were recorded in another six isolates. The most salient example was 1987 (Table 1), a strain hosting 10 viruses belonging to four taxonomic groups. The highest documented natural virus load in *Heterobasidion* consisted of eight dsRNA or ssRNA viruses in a single *H. parviporum* strain (Sutela et al. 2021), whereas in a transmission experiment, the highest achieved number of coinfecting dsRNA viruses was five (Hantula et al. 2020). Infections with comparably high numbers of related and unrelated viruses are known in other fungi (Mu et al. 2021; Li et al. 2023) and oomycetes (Botella et al. 2022; Raco et al. 2022).

**Table 1.** Mycoviruses detected.

| Mycovirus | Acronym | Segment | GenBank accession | Host strain | Length (nt) | % GC | Mapping reads<sup>h, i</sup> | Mean depth<sup>i</sup> | BLASTX first hit by total score<sup>j</sup> | Identity<sup>l</sup> | Query cover<sup>k</sup> | E value |
| --- | --- | --- | --- | --- | --- | --- | --- | --- | --- | --- | --- | --- |
| Heterobasidion mitovirus 3 | HetMV3 | RNA1 | ON014534 | *H. annosum* 2009 | 4933 | 43.1% | 228,580 | 6850 | Heterobasidion mitovirus 3 (QED55407)<sup>k</sup> | 96.9%<sup>l</sup> | 49%<sup>k</sup> | 0<sup>k</sup> |
| Heterobasidion narna-like virus 2 | HetNIV2 | RNA1 | ON014535 | *H. annosum* 1987<sup>a</sup> | 3877 | 53.7% | 57,684/<br>51,735 | 2187/<br>1919 | Heterobasidion narna-like virus 1 (UHK02569) | 80.3% | 94% | 0 |
| Heterobasidion narna-like virus 4 | HetNIV4 | RNA1 | ON985234 | *H. annosum* 1987<sup>a</sup> | 3970 | 54.0% | 62,816/<br>42,325 | 2329/<br>1538 | Heterobasidion narna-like virus 1 (UHK02569) | 80.2% | 95% | 0 |
|  |  | RNA2 | ON014536 |  | 4011 | 55.5% | 40,456/<br>150,103 | 1484/<br>5410 | Heterobasidion narna-like virus 1 (UHK02570) | 60.5% | 87% | 0 |
| Heterobasidion ourmia-like virus 2 | HetOIV2 | RNA1 | ON014537 | *H. annosum* 2072 | 2783 | 57.4% | 438 | 23 | Leucocoprinus ourmiavirus B (QED42953) | 51.8% | 46% | 8e−111 |
| Heterobasidion ourmia-like virus 3 | HetOIV3 | RNA1 | ON014538 | *H. annosum* 2038<sup>b</sup> | 2632 | 45.1% | 5122 | 286 | Heterobasidion ourmia-like virus 1 (UHK02571) | 48.6% | 52% | 7e−110 |
| Heterobasidion fusarivirus 1 | HetFV1 | RNA1 | ON014533 | *H. annosum* 1987 | 6913 | 54.4% | 444/<br>2101 | 9/<br>43 | Heterobasidion parviporum fusarivirus 1 (WOK44148) | 91.5% | 82% | 0 |
| Heterobasidion tobamo-like virus 1 | HetTIV1 | RNA1 | ON014539 | *H. annosum* 1989 | 12,586 | 46.9% | 1785 | 20 | Lentinula edodes tobamo-like virus 1 (QOX06054) | 35.9% | 29% | 1e−66 |
| Heterobasidion ambi-like virus 3 | HetAIV3 | RNA1 | ON014526 | *H. parviporum* 1991 | 4947 | 48.7% | 740 | 22 | Heterobasidion ambi-like virus 3 (UHK02576) | 95.8% | 42% | 0 |
| Heterobasidion ambi-like virus 5 | HetAIV5 | RNA1 | ON014527 | *H. annosum* 2050 | 4815 | 47.7% | 5461 | 166 | Heterobasidion ambi-like virus 1 (UHK02572) | 76.9% | 37% | 0 |
| Heterobasidion ambi-like virus 6 | HetAIV6 | RNA1 | ON014528 | *H. annosum* 2028<sup>c</sup> | 5108 | 46.8% | 9624 | 276 | Armillaria novae-zelandiae ambi-like virus 1 (DAD54841) | 32.9% | 40% | 4e−94 |
| Heterobasidion ambi-like virus 7 | HetAIV7 | RNA1 | ON014529 | *H. annosum* 2040<sup>d</sup> | 4965 | 47.1% | 8050 | 236 | Heterobasidion ambi-like virus 4 (UHK02578) | 57.2% | 41% | 0 |
| Heterobasidion ambi-like virus 8 | HetAIV8 | RNA1 | ON014530 | *H. annosum* 1983<sup>e</sup> | 5026 | 47.5% | 8346 | 243 | Heterobasidion ambi-like virus 4 (UHK02578) | 33.8% | 39% | 2e−90 |
| Heterobasidion<br>ambi-like virus 9 | HetAlV9 | RNA1 | ON014531 | *H. annosum*<br>2028 | 4903 | 48.5% | 1140 | 34 | Heterobasidion<br>ambi-like virus 3 (UHK02576) | 64.2% | 42% | 0 |
| Heterobasidion<br>ambi-like virus 19 | HetAlV19 | RNA1 | ON985229 | *H. annosum*<br>1987 | 4977 | 47.3% | 8922/<br>18,873 | 260/<br>540 | Heterobasidion<br>ambi-like virus 4 (UHK02578) | 56.5% | 41% | 0 |
| Heterobasidion<br>ambi-like virus 20 | HetAlV20 | RNA1 | ON985230 | *H. annosum*<br>1987 | 4919 | 48.9% | 7517/<br>47,340 | 225/<br>1389 | Heterobasidion<br>ambi-like virus 3 (UHK02577) | 63.9% | 40% | 0 |
| Heterobasidion<br>ambi-like virus 21 | HetAlV21 | RNA1 | ON985231 | *H. annosum*<br>1987 | 4862 | 48.0% | 199,680/<br>1,487,296 | 6006/<br>44,221 | Heterobasidion<br>ambi-like virus 1 (UHK02572) | 78.0% | 37% | 0 |
| Heterobasidion<br>ambi-like virus 22 | HetAlV22 | RNA1 | ON985232 | *H. annosum*<br>1987 | 4939 | 47.4% | 26,585/<br>185,342 | 784/<br>5381 | Heterobasidion<br>ambi-like virus 4 (UHK02578) | 60.2% | 41% | 0 |
| Heterobasidion<br>ambi-like virus 23 | HetAlV23 | RNA1 | ON985233 | *H. annosum*<br>1987 | 4882 | 48.3% | 397/<br>454 | 12/<br>13 | Heterobasidion<br>ambi-like virus 4 (UHK02578) | 47.4% | 41% | 0 |
| Heterobasidion<br>ambi-like virus 24 | HetAlV24 | RNA1 | OR031238 | *H. annosum*<br>1993 | 4995 | 46.9% | 8227 | 237 | Heterobasidion<br>ambi-like virus 4 (UHK02578) | 61.8% | 41% | 0 |
| Heterobasidion<br>ambi-like virus 25 | HetAlV25 | RNA1 | OR031239 | *H. annosum*<br>1987 | 5027 | 46.9% | 6483/<br>19,649 | 191/<br>561 | Heterobasidion<br>ambi-like virus 4 (UHK02578) | 56.7% | 41% | 0 |
| Heterobasidion<br>ambi-like virus 26 | HetAlV26 | RNA1 | OR031240 | *H. annosum*<br>1983 | 5005 | 46.4% | 14,219 | 411 | Heterobasidion<br>ambi-like virus 4 (UHK02578) | 54.3% | 41% | 0 |
| Heterobasidion<br>ambi-like virus 27 | HetAlV27 | RNA1 | OR031241 | *H. annosum*<br>1983 | 4914 | 49.0% | 6144 | 184 | Heterobasidion<br>ambi-like virus 3 (UHK02577) | 61.7% | 40% | 0 |
| Heterobasidion<br>ambi-like virus 28 | HetAlV28 | RNA1 | OR031242 | *H. annosum*<br>1993 | 2537<sup>f</sup> | 47.4% | 4143 | 219 | Heterobasidion<br>ambi-like virus 4 (UHK02579) | 46.0% | 77% | 0 |
| Heterobasidion<br>ambi-like virus 29 | HetAlV29 | RNA1 | OR031243 | *H. annosum*<br>2028 | 2390<sup>f</sup> | 45.9% | 1699 | 102 | Heterobasidion<br>ambi-like virus 4 (UHK02578) | 61.8% | 42% | 0 |
| Coguvirus movement<br>protein-like sequence | CVMPIS | RNA2 | ON014532 | *H. annosum*<br>1987 | 2425<sup>g</sup> | 55.5%<sup>g</sup> | 578<sup>g</sup>/<br>2666<sup>g</sup> | 35<sup>g</sup>/<br>156<sup>g</sup> | Jiangsu sediment phenui-like<br>virus (QYF49545) | 24.9% | 24% | 4e-04 |

<sup>a</sup> Other *H. annosum* hosts were 1983 and 1993.
<sup>b</sup> Other *H. annosum* hosts were 1887 and 2028.
<sup>c</sup> Other *H. annosum* hosts were 1983, 1993, 2038 and 2050.
<sup>d</sup> Another *H. annosum* host was 2028.
<sup>e</sup> Another *H. annosum* host was 1993.
<sup>f</sup> Partial genomic sequence.
<sup>g</sup> Without polyT tail.
<sup>h</sup> Using Geneious assembler with medium-low sensitivity.
<sup>i</sup> A single number pertains to the pooled library in which the virus was first detected and the host strain belongs to; if there are two numbers, the first one pertains to library ANNO1 and the second one to library 1987.

### 3.2. Mitovirus

A new strain of Heterobasidion mitovirus 3 (HetMV3) was detected, sharing 97% BLASTX identity with the strain HetMV3-an1 (Vainio, 2019; Table 1). Both strains inhabited *H. annosum*, but the virus was also recorded in *H. parviporum* (Vainio 2019; Sutela et al. 2021). The nucleotide (nt) sequence identities shared by the Czech HetMV3 strain and other strains originating from Poland, Finland and Russia (Vainio 2019) were 94%, 90% and 90%, respectively. This indicates intraspecies genomic differentiation with increasing geographic distance, already noted by Vainio (2019). The HetMV3 genome contains the mitoviral RdRp conserved domain (Fig. 2). Phylogenetically, it clustered together with HetMV1 within the *Duamitovirus* genus of the *Mitoviridae* family (Fig. 3), confirming the analysis of Vainio (2019). Despite infecting a single host strain, HetMV3 had the highest read coverage among all viruses found in our study, a mean depth of 6850 (Table 1). Outstanding accumulation levels of HetMV3 strains relative to other viruses detected in the same RNA-Seq library have previously been observed also in *H. parviporum* (Sutela et al. 2021). The high accumulation levels of Rhizoctonia cerealis duamitoviruses in their host compared with other viruses have been explained by the protective environment of mitochondria against cytoplasmic RNA interference (RNAi) (Li et al. 2023), which was also demonstrated for Cryphonectria parasitica mitovirus 1 (Shahi et al. 2019). Our data supports the notion that the mitochondrial replication of mitoviruses is a successful strategy to avoid RNA silencing, even though the RNAi machinery of *Heterobasidion* was shown to affect HetMV1 to a certain degree (Vainio et al. 2015a).

**Fig. 2.**
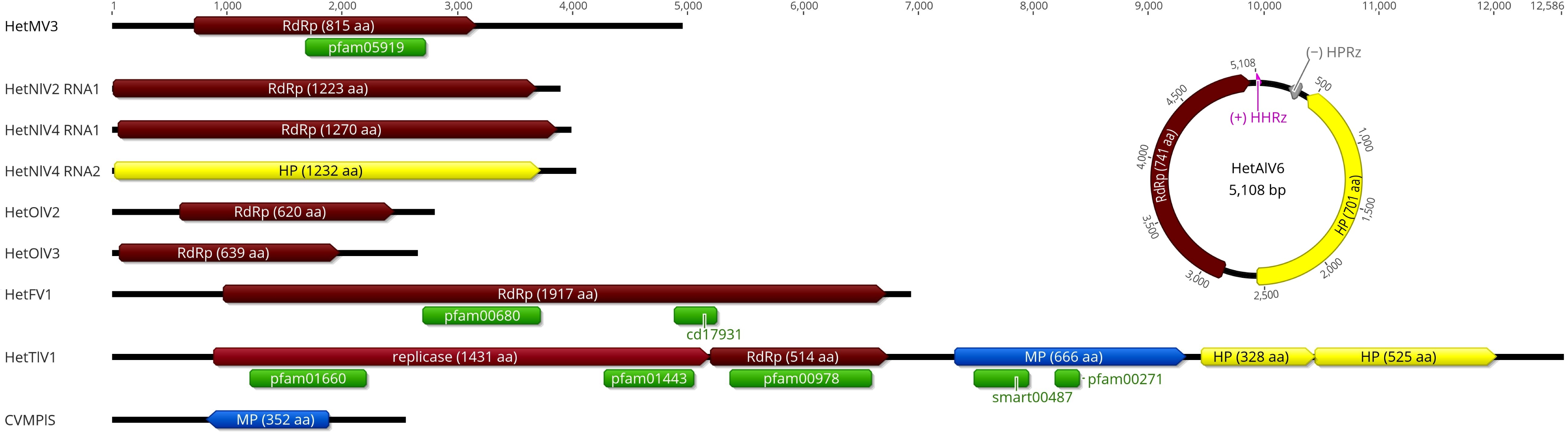
Genome organization of mycoviruses detected, drawn to scale. One representative of the novel ambi-like viruses was selected to illustrate the typical location of ribozymes within their circular genome. HPRz, hairpin ribozyme; HHRz, hammerhead ribozyme. Predicted open reading frames: RdRp, RNA-dependent RNA polymerase; HP, hypothetical protein; MP, movement protein. Conserved domains: pfam05919, mitovirus RdRp (E value: 9.17e−84); pfam00680, RdRp (1.23e−15); cd17931, DEXH-box helicase (5.37e−08); pfam01660, methyltransferase (8.56e−14); pfam01443, helicase (4.00e−29); pfam00978, RdRp (4.65e−78); smart00487, DEAD-like helicases superfamily (2.83e−06); pfam00271, helicase conserved C-terminal domain (5.78e−05)

**Fig. 3.**
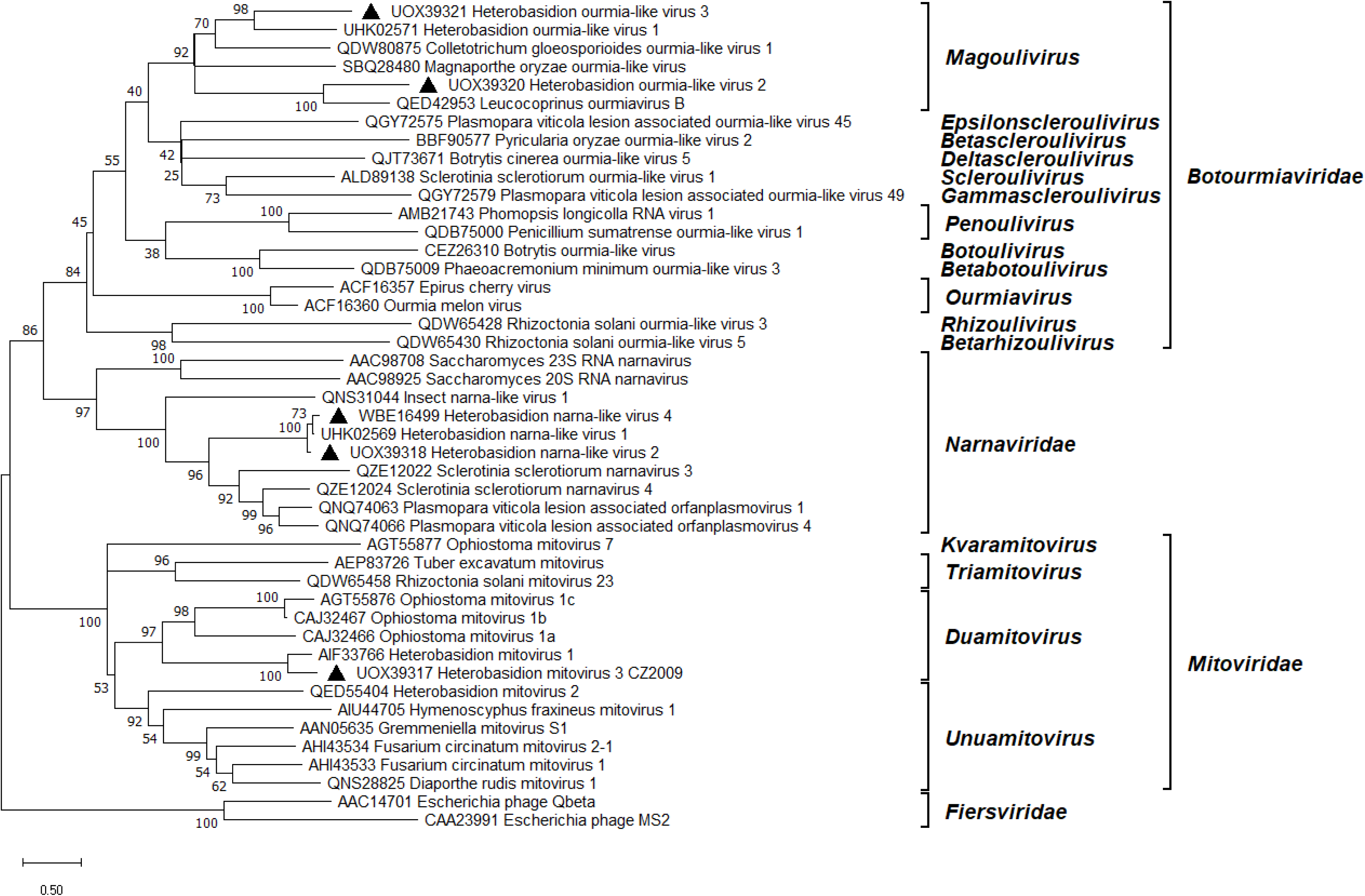
Phylogenetic relationships within and between relevant families of the *Lenarviricota* phylum. The *Fiersviridae* family served as outgroup. The tree was built based on the alignment of RdRp aa sequences generated with MUSCLE in MEGA11 (Tamura et al. 2021). The evolutionary history was inferred by using the Maximum Likelihood method and Whelan And Goldman + Freq. model (Whelan and Goldman 2001). A discrete Gamma distribution was used to model evolutionary rate differences among sites (five categories +*G* +*I*). All positions with less than 95% site coverage were eliminated, i.e., fewer than 5% alignment gaps, missing data, and ambiguous bases were allowed at any position. Evolutionary analyses were conducted in MEGA11 with 500 bootstrap repeats. The percentage of trees in which the associated taxa clustered together is shown next to the branches. Branch lengths are proportional to the number of substitutions per site. Ourmia- and narna-like viruses and the mitovirus strain described in this study are denoted by a triangle

### 3.3. Narna-like viruses

Two contigs resembling the RNA1 segment of Heterobasidion narna-like virus 1 (HetNlV1) (Sutela et al. 2021) were found, each with 80% identity in BLASTX (Table 1). Another contig shared 61% aa identity with the hypothetical protein (HP) encoded by the HetNlV1 RNA2 segment. Similarly to HetNlV1, the length of each detected virus segment was ca. 4 kilobases (kb). All three segments had a high read coverage in datasets ANNO1 and 1987 (Table 1), and their presence was confirmed in the same three isolates. The hyphal tip isolation experiment clarified that two of the segments belong to the same virus, being always cotransmitted into the same monohyphal cultures of 1987 with a transmission frequency of 90%, as opposed to the third segment’s rate of 20%. Based on these results, the unmatched RNA1 segment was designated as HetNlV2, while a new bisegmented narna-like virus was named HetNlV4 (Table 1; Fig. 2). The HetNlV4 RNA1 segment would have remained undetected based on analyzing only the ANNO1 library. The two viruses shared 75% aa identity over their RdRp. Pw identities of RT-PCR amplicons of narna-like virus strains ranged 94–99% at both the nt and aa levels (Supplemental Fig. S2). *Heterobasidion* narna-like viruses formed a highly supported cluster within *Narnaviridae* (Fig. 3). Their closest relatives were the multisegmented narna-like viruses of *S. sclerotiorum* (Jia et al. 2021) and the bipartite Plasmopara viticola lesion associated orfanplasmoviruses (Chiapello et al. 2020). HetNlV2 and HetNlV4 always occurred in coinfection with at least four other viruses, most frequently ambi-like viruses (Table 1), which was also the case for HetNlV1 infecting *H. parviporum* (Sutela et al. 2021).

### 3.4. Ourmia-like viruses

Two distinct ourmia-like viruses were discovered. For one of them, the most significant BLASTX hit was Leucocoprinus ourmiavirus B with 52% identity, while the other one showed highest similarity (49% BLASTX identity) to Heterobasidion ourmia-like virus 1 (HetOlV1) (Sutela et al. 2021). We named them HetOlV2 and HetOlV3, respectively (Table 1). Their nonsegmented genomes of 2.8 and 2.6 kb contain a single predicted ORF, encoding for a putative RdRp (Fig. 2). HetOlV2 dwelled in a single isolate and had a very low coverage, a mean depth of 23 reads. HetOlV3 was confirmed to infect three host isolates and reached moderate accumulation levels in the ANNO2 dataset (Table 1). Pw identities of RT-PCR amplicons of HetOlV3 strains ranged 89–91% at the nt level and 91–95% at the aa level (Supplemental Fig. S2). Phylogenetic analysis of the *Lenarviricota* phylum showed that *Heterobasidion* ourmia-like viruses resemble members of the *Magoulivirus* genus within *Botourmiaviridae* (Fig. 3). HetOlV3 was responsible for single or multiple infections with up to four ambi-like viruses (Table 1). In comparison, HetOlV1 in *H. parviporum* had a propensity for coinfections with various dsRNA and ssRNA viruses (Sutela et al. 2021).

### 3.5. Fusarivirus

We report the detection of a fusarivirus as a novelty in the virome of *Heterobasidion* spp. The relevant contig assembled from the ANNO1 dataset was 3.3 kb long, representing only a partial genome lacking both ends of an ORF. Data from 1987 yielded the full genome of Heterobasidion fusarivirus 1 (HetFV1), which had the lowest read coverage among the viruses detected in this study (Table 1). HetFV1 gave highly significant matches with members of the *Fusariviridae* family. The first BLASTX hit was the unpublished sequence of Heterobasidion parviporum fusarivirus 1 (HetpaFV1) from Finland (GenBank OR644501) with 92% identity. The HetFV1 genome sequence of nearly 7 kb contains one ORF comprising an RdRp conserved domain and a DEXH-box helicase domain (Fig. 2). The aa sequence motifs within the RdRp were highly conserved among selected fusariviruses; HetFV1 shared the most similarities with Phlebiopsis gigantea fusarivirus 1 (Supplemental Fig. S3). As *Heterobasidion* spp. and *P. gigantea* often share the same ecological niche, conifer stumps, this observation brings up the question whether the horizontal transmission of the common ancestor of these fusariviruses could have occurred between the two fungal taxa. Within the helicase domain, the DEXH motif of HetFV1 was identical to that of certain selected members of *Fusariviridae* and *Flaviviridae* (Supplemental Fig. S4). Phylogenetically, HetFV1 clustered in the *Gammafusarivirus* genus with HetpaFV1 and fusariviruses of *Lentinula edodes* (Guo et al. 2021) and *P. gigantea* (Drenkhan et al., 2022; Fig. 4).

**Fig. 4.**
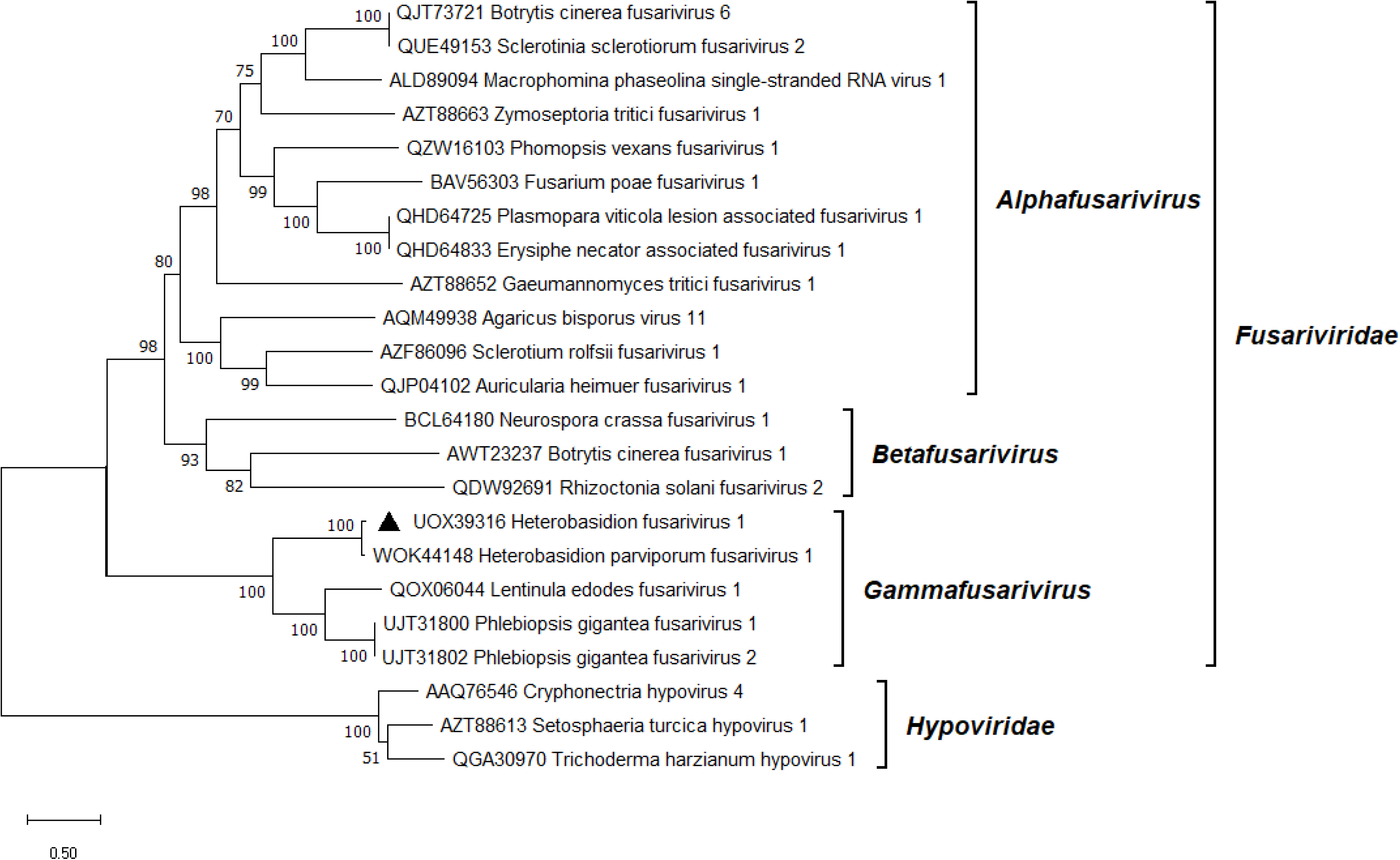
Phylogenetic relationships of HetFV1 (denoted by a triangle) with selected members of the *Fusariviridae* family. The *Hypoviridae* family served as outgroup. The tree was built based on the alignment of RdRp aa sequences generated with MUSCLE in MEGA11 (Tamura et al. 2021). The evolutionary history was inferred by using the Maximum Likelihood method and Le_Gascuel_2008 model (Le and Gascuel 2008). A discrete Gamma distribution was used to model evolutionary rate differences among sites (five categories +*G* +*I*). All positions with less than 95% site coverage were eliminated, i.e., fewer than 5% alignment gaps, missing data, and ambiguous bases were allowed at any position. Evolutionary analyses were conducted in MEGA11 with 1000 bootstrap repeats. The percentage of trees in which the associated taxa clustered together is shown next to the branches. Branch lengths are proportional to the number of substitutions per site

### 3.6. Tobamo-like virus

A contig showing moderate similarity to members of the *Virgaviridae* family, with Lentinula edodes tobamo-like virus 1 (Guo et al. 2021) as its first hit in BLASTX, was named Heterobasidion tobamo-like virus 1 (HetTlV1). It caused a low titer infection of a single host strain, the mean depth of coverage was merely 20 reads (Table 1). The first tobamo-like virus in *Heterobasidion* is monopartite, 12.6 kb long (Fig. 2). Its largest ORF was predicted to encode a replicase including methyltransferase and helicase domains. Varying degrees of conservation of aa sequence motifs were observed across *Virgaviridae*, but overall, HetTlV1 showed highest similarity to Podosphaera prunicola tobamo-like virus within both domains (Supplemental Figs. S5 and S6). The second ORF encodes the RdRp, which appeared slightly more conserved than the upstream regions. Taking into account all aa sequence motifs within the RdRp, HetTlV1 was most similar to Plasmopara viticola lesion associated tobamo-like virus 1 (Supplemental Fig. S7). The third ORF resembles a MP with DEAD-like helicases superfamily and helicase conserved C-terminal domains (Fig. 2). The last two ORFs are of unknown function. The number of nt between the ORFs encoding the replicase and the RdRp suggests that −1 ribosomal frameshifting is utilized to express these proteins. Similarly, +1 ribosomal frameshift appears to take place between the last two ORFs encoding HPs. The mechanism behind these frameshifts is unclear as neither typical shifty heptamer motifs nor octanucleotides promoting programmed ribosomal frameshifting (Firth and Brierley 2012) were noticed at the ends of the upstream ORFs. However, the first 200 nt of the downstream ORFs contain the UUAUG sequence, which is deemed to be a key motif for translation (Raco et al. 2022). Programmed −1 ribosomal frameshift has so far been described in one tobamovirus species, *Hibiscus latent Singapore virus*, whose hepta-adenosine stretch located in its replicase gene triggers the frameshift (Niu et al. 2014). Phylogenetic analysis placed HetTlV1 as a separate individual branch at a large distance from classified and unclassified members of *Virgaviridae* (Fig. 5).

**Fig. 5.**
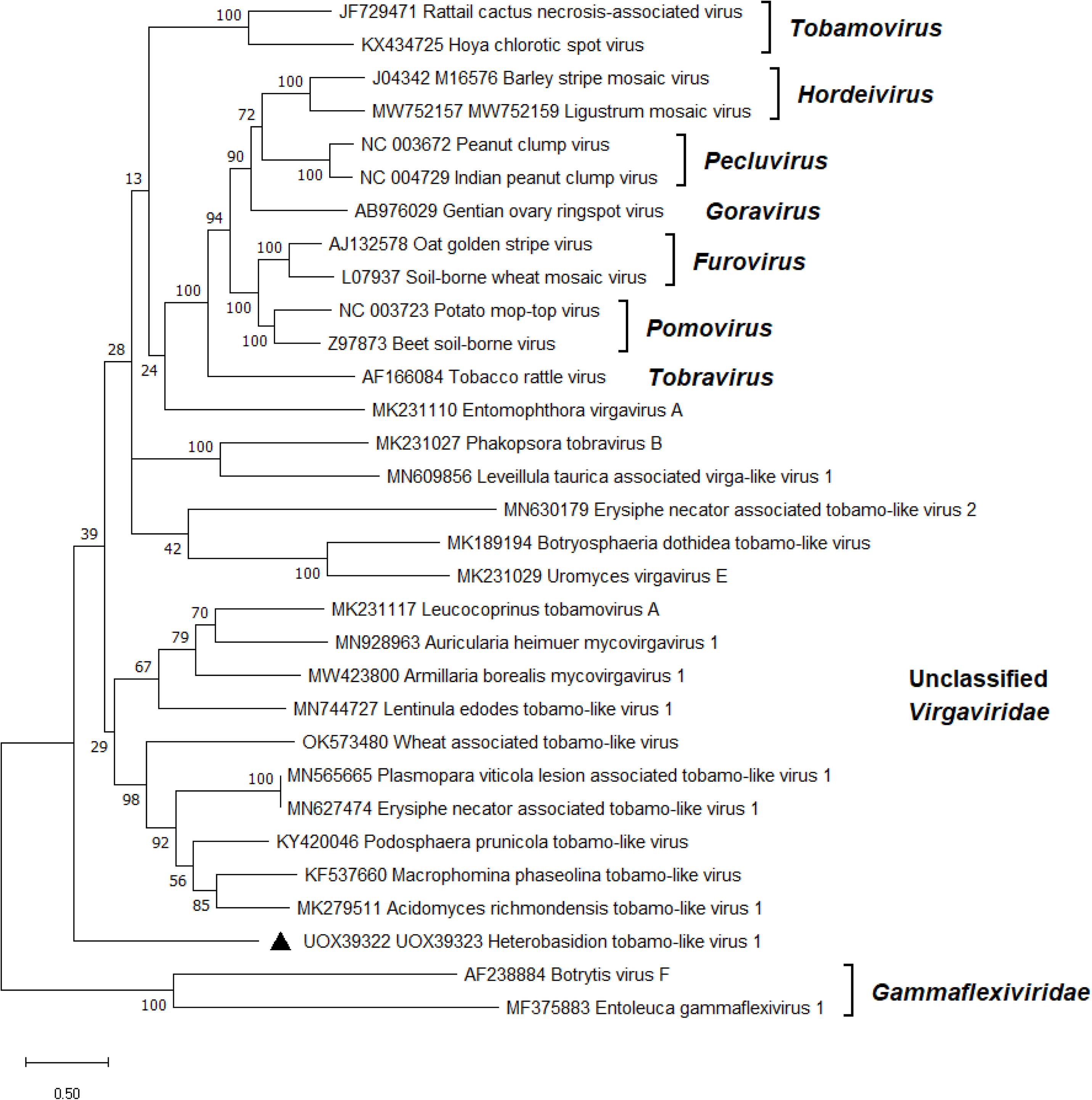
Phylogenetic relationships of HetTlV1 (denoted by a triangle) with selected members of the *Virgaviridae* family. The *Gammaflexiviridae* family served as outgroup. The tree was built based on the codon-aligned nt sequences of the replication proteins and RdRps generated with MUSCLE in MEGA11 (Tamura et al. 2021). The evolutionary history was inferred by using the Maximum Likelihood method and Kimura 2-parameter model (Kimura 1980). A discrete Gamma distribution was used to model evolutionary rate differences among sites (five categories +*G* +*I*). All positions with less than 95% site coverage were eliminated, i.e., fewer than 5% alignment gaps, missing data, and ambiguous bases were allowed at any position. Evolutionary analyses were conducted in MEGA11 with 500 bootstrap repeats. The percentage of trees in which the associated taxa clustered together is shown next to the branches. Branch lengths are proportional to the number of substitutions per site

### 3.7. Ambi-like viruses

Ambiviruses constitute a recently discovered but rapidly expanding group of mycoviruses, proposed to be classified as a new phylum by the International Committee on Taxonomy of Viruses (ICTV) (Turina et al., 2023). Their distinctive features include the possession of two non-overlapping ORFs in ambisense orientation (Sutela et al. 2020) and the activity of self-cleaving ribozymes enabling rolling circle replication (Forgia et al. 2023). The remarkable abundance of ambi-like viruses has been reported via data mining in metatranscriptomes (Forgia et al. 2023; Lee et al. 2023) and fungal transcriptomes (Chong and Lauber 2023). The present study was no exception. One of the detected contigs shared 96% BLASTX identity with the virus strain HetAlV3-pa1 known from Finland (Sutela et al., 2021; Table 1). The 3’ terminus of the HetAlV3 genome sequence was completed by RACE based on three cloned amplicons spanning 1269 bp. Notably, the read coverage of HetAlV3 was markedly low both in the study by Sutela et al. (2021) and in the present work. Apart from HetAlV3, 10 new ambi-like viruses were retrieved from our pooled datasets, designated as HetAlV5–9, HetAlV24 and HetAlV26–29 (Table 1). The RNA-Seq analysis of *H. annosum* strain 1987 revealed the existence of six further ambi-like viruses, named HetAlV19–23 and HetAlV25 (Table 1). These originally went undetected in the ANNO1 dataset, but their genome sequences were posteriorly shown to have varying levels of read support in both datasets (Table 1). A second RT-PCR was needed to obtain a well visible amplicon within the RdRp encoding ORF of HetAlV23 and HetAlV29, both of which had low read coverage, as well as that of HetAlV6 from isolate 2038. Most likely, HetAlV6 caused a low titer infection in this particular isolate as it showed a moderate read coverage while being present in five *H. annosum* strains altogether (Table 1). According to BLASTX search, the newly discovered ambi-like viruses were 33–78% identical to known ambi-like viruses of *Heterobasidion* or *Armillaria* (Table 1). As usual for ambiviruses, each virus reported here had a genome of approximately 5 kb, comprising two bidirectional ORFs. The contigs resembling ambi-like viruses were almost always assembled with the putative RdRp-encoding ORF in the sense orientation. Contigs assembled in the reverse or both orientations were reverse complemented, and the final genomic sequences were deposited into GenBank with the putative RdRp in the coding orientation. In several cases, the ambiviral contigs were assembled as dimers or trimers of the same genomic sequence. This observation is consistent with previous studies (Sutela et al. 2020; Forgia et al. 2021), indicating that ambiviruses have circular genomes (Turina et al. 2023). Each ambi-like virus contained at least one hairpin (HPRz) or hammerhead (HHRz) ribozyme, typically positioned at the C-termini of both ORFs (Supplemental Table S3). The genome organization of HetAlV6, representative of the novel ambi-like viruses is depicted in Fig. 2. For HetAlV28 and HetAlV29, only a partial genome was determined, lacking either the RdRp (ORFA) or the HP (ORFB). No conserved domains were found, but the longest predicted protein of each virus with a complete genome included the GDD motif, which is considered the hallmark of RdRps. Virus candidates showing >90% nt pw identity over their putative RdRp encoding region were considered strains of the same virus species. This is an arbitrary threshold/species demarcation criterion suggested by the ICTV experts based upon existing data (Turina et al. 2023). Pw identities of RT-PCR amplicons of ambi-like virus strains ranged 88–97% at the nt level and 95–100% at the aa level (Supplemental Fig. S2). The similarity of RdRps was generally slightly higher than that of HPs. The ambi-like viruses of *Heterobasidion* are presumably of polyphyletic origin as they were placed in three separate branches (Fig. 6). In two bigger clusters, they grouped with each other, whereas HetAlV6 was more closely associated with Armillaria novae-zelandiae ambi-like virus 1 (Linnakoski et al. 2021). With two exceptions (isolates 1991 and 2040), the *Heterobasidion* ambi-like viruses caused coinfections in their host strains with CVMPlS, HetFV1, ambi-, narna- or ourmia-like viruses (Table 1). HetAlV1–4 were always found in mixed infections in Finnish *H. parviporum* isolates (Sutela et al. 2021). Ambi-like viruses of other basidiomycetes, such as *Tulasnella* spp. (Sutela et al. 2020), *Armillaria* spp. (Linnakoski et al. 2021) and *P. gigantea* (Drenkhan et al. 2022) were shown to be responsible for both single and multiple virus infections. On the other hand, Ceratobasidium ambivirus 1 (Sutela et al. 2020) and Cryphonectria parasitica ambivirus 1 (Forgia et al. 2021) were detected in single infections.

**Fig. 6.**
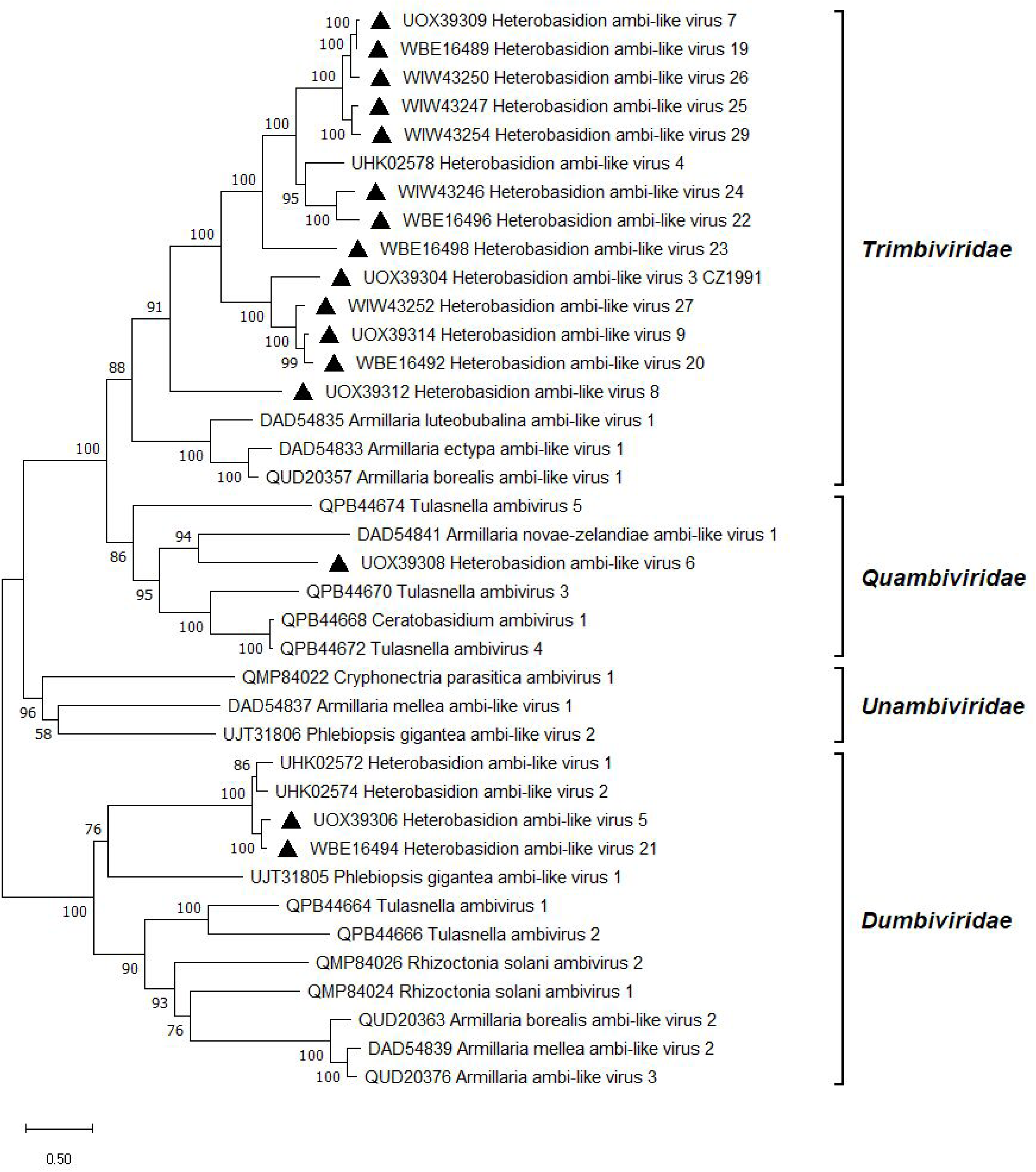
Phylogenetic relationships of ambi-like viruses. The tree was built based on the alignment of RdRp aa sequences generated with MUSCLE in MEGA11 (Tamura et al. 2021). The evolutionary history was inferred by using the Maximum Likelihood method and Le_Gascuel_2008 model (Le and Gascuel 2008). A discrete Gamma distribution was used to model evolutionary rate differences among sites (five categories +*G* +*I*). All positions with less than 95% site coverage were eliminated, i.e., fewer than 5% alignment gaps, missing data, and ambiguous bases were allowed at any position. Evolutionary analyses were conducted in MEGA11 with 1000 bootstrap repeats. The percentage of trees in which the associated taxa clustered together is shown next to the branches. Branch lengths are proportional to the number of substitutions per site. Viruses described in this study are denoted by a triangle

### 3.8. Coguvirus movement protein-like sequence

The cDNA libraries ANNO1 and 1987 yielded a contig with resemblance to the MP encoding segment of plant coguviruses. The contig was most similar (25% identity in BLASTX) to the putative MP of the unclassified Jiangsu sediment phenui-like virus (Chen et al. 2022). Putative segments encoding for the polymerase and the nucleocapsid protein were absent in the data, only a 2.5 kb long segment of the positive-sense antigenome was assembled. The genomic segment encompasses a polyT tail of 109 nt and encodes a MP (Fig. 2). DNA analysis indicated that the sequence was not genome integrated. The fragment was designated as Coguvirus movement protein-like sequence (CVMPlS; Table 1) and represents the first putative negative-strand RNA virus in *Heterobasidion*. CVMPlS seems to be distantly related to classified members of the *Phenuiviridae* family (Fig. 7), although a tree built on the MP sequence may only give an approximate idea of its phylogenetic position. CVMPlS infected the multivirus isolate 1987 together with a fusarivirus, ambi- and narna-like viruses (Table 1).

**Fig. 7.**
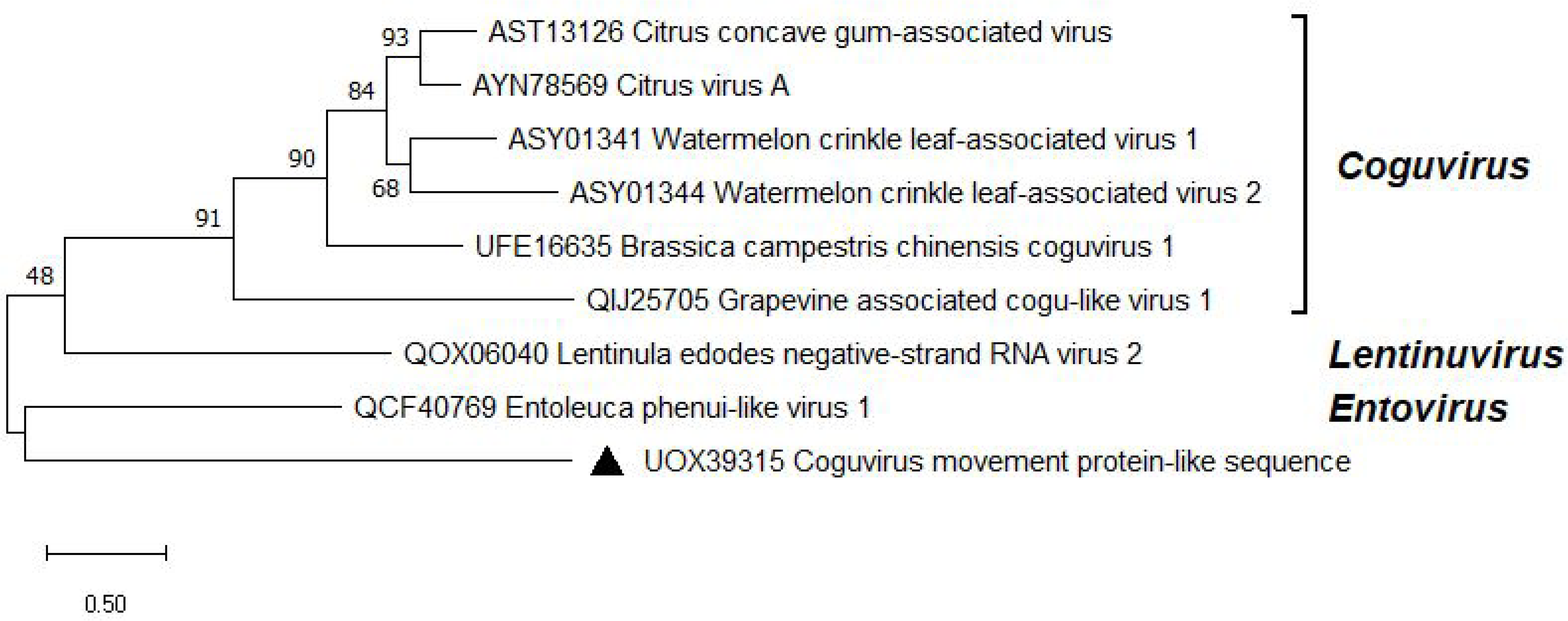
Phylogenetic relationships of CVMPlS (denoted by a triangle) with selected members of the *Phenuiviridae* family. The tree was built based on the alignment of MP aa sequences generated with MUSCLE in MEGA11 (Tamura et al. 2021). The evolutionary history was inferred by using the Maximum Likelihood method and Le_Gascuel_2008 model (Le and Gascuel 2008). A discrete Gamma distribution was used to model evolutionary rate differences among sites (five categories +*G*). All positions with less than 95% site coverage were eliminated, i.e., fewer than 5% alignment gaps, missing data, and ambiguous bases were allowed at any position. Evolutionary analyses were conducted in MEGA11 with 1000 bootstrap repeats. The percentage of trees in which the associated taxa clustered together is shown next to the branches. Branch lengths are proportional to the number of substitutions per site

### 3.9. Efficiency of virus elimination and horizontal transmission

By the time we commenced the virus removal (curing) experiments, HetAlV23 had already been lost from isolate 1987 during its storage at 4°C for 1 year. This could be explained by its extremely low accumulation level, a mean depth of 12 reads (Table 1). Viruses were retained in the monohyphal cultures in widely varying frequencies (0–100%). HetTlV1 was absent from the hyphal tip isolates of strain 1989. Contrastingly, HetOlV2 was transmitted to 10 out of 10 monohyphal cultures of strain 2072. In strain 1987, which was infected by multiple viruses, the success rates of virus elimination were more nuanced. HetAlV20–22 were retained in all cases, HetNlV4 in 90%, HetFV1 in 70%, HetNlV2 in 20% of the monohyphal cultures, while HetAlV19, HetAlV25 and CVMPlS were absent. Generally, viruses perceived to cause infections of low titer based on their read coverage were less stable. Still, characteristics of the virus and/or the fungal genotype also seemed to influence the likelihood of transmission. These findings are consistent with the results of Ihrmark et al. (2002), who found that the higher concentration of dsRNA in an *H. parviporum* isolate increased the dsRNA transmission rate into single hyphal tip isolates.

The original isolates 1987, 1989 and 2072, as well as the monohyphal cultures 1987–A2, – A4, –A5 and –C1, containing different combinations of viruses (Fig. 8A), were fully cured of their viral infections after both 1 and 2 weeks of thermal treatment. Temperature treatment lasting 2 or 4 weeks at 32–33°C was previously successfully applied to cure *Heterobasidion* strains of partitiviruses (Vainio et al. 2010; Vainio et al. 2018), but even lower temperatures have been shown efficient: a two-week incubation at 28°C resulted in loss of the dsRNA genomes of partitiviruses and curvulaviruses in *H. parviporum* (Jalo 2008).

**Fig. 8.**
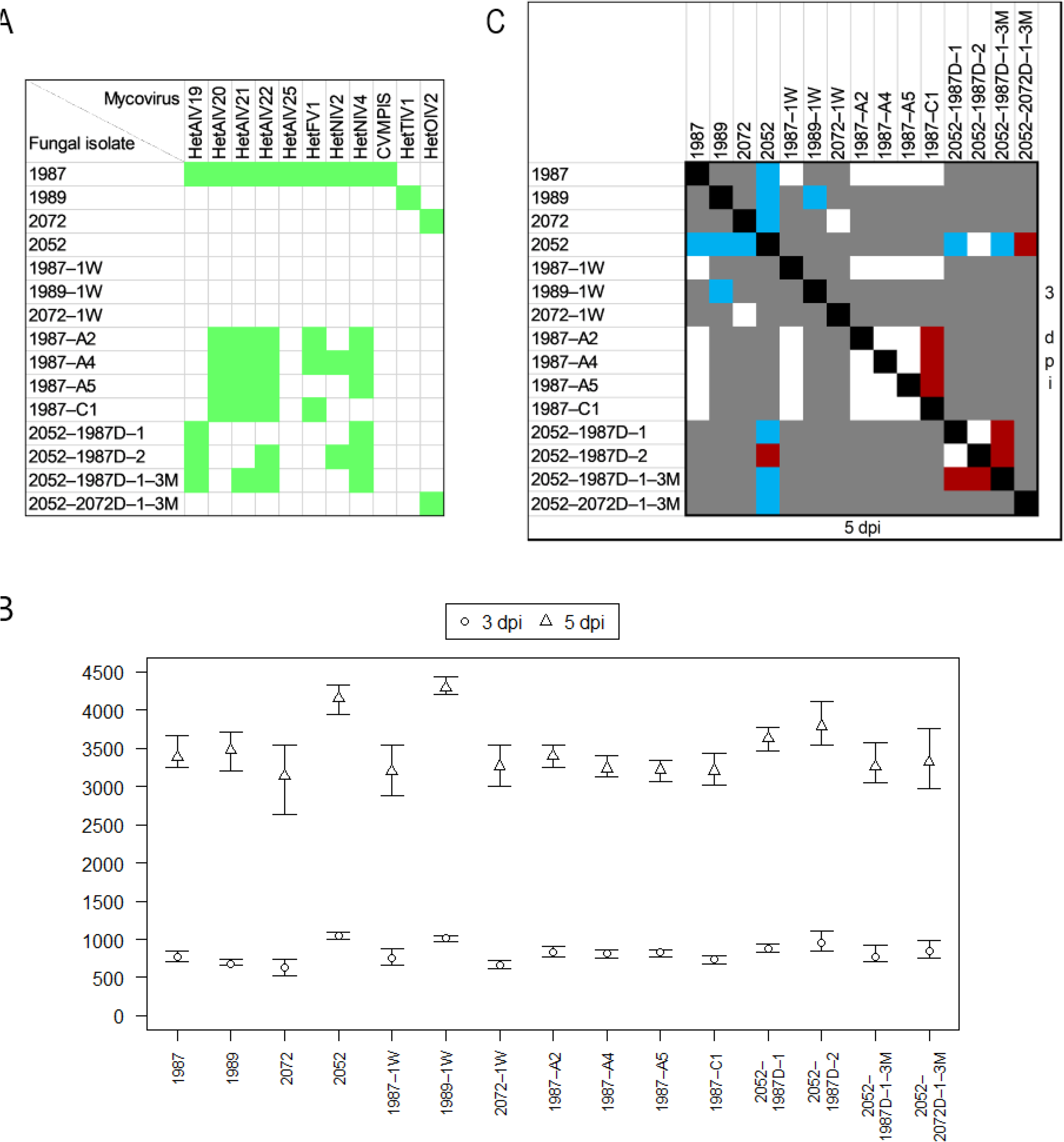
A. Presence of viruses in *H. annosum* isolates used for the growth rate experiment. Note that HetAlV23 had been lost from 1987 during its storage at 4°C for 1 year. –1W, isogenic isolates fully cured of viral infections by thermal treatment lasting 1 week; –A2, – A4, –A5, –C1, monohyphal cultures created by hyphal tip isolation; 2052–, isogenic isolates, which have received different combinations of viruses by horizontal transmission from a donor strain denoted by D; –3M, dual cultures were maintained for 3 months (where absent, the isolation was performed after 1 month). **B** Mean area [mm^2^] of 2% malt extract agar plates covered by mycelium 3- and 5-days post-inoculation (dpi). The bars represent the range of values of five technical replicates. **C** Statistical significance of differences in the growth rate between pairs of isolates 3 and 5 dpi, calculated with *t*-test. Blue, *p*<0.001; red, *p*<0.05; white, *p*>0.05; gray, not tested

After the maintenance of dual cultures for 1 month, horizontal transmission of virus(es) occurred in just one out of nine pairings. Isolate 2052 received HetAlV19 and HetNlV4 from the donor strain 1987 when sampled from spot 1 (Figs. 1A and 8A). Interestingly, two additional viruses, HetAlV22 and HetNlV2 were transmitted to 2052 when sampled from spot 2, which was located farther from the interaction zone (Figs. 1A and 8A). Three months post-inoculation, 2052 received the same four viruses from 1987 as 1 month post-inoculation when sampled from spots 2 and 3 (Fig. 1B). However, the sample from spot 1 was infected by a different combination of viruses, i.e. HetAlV21 replaced HetNlV2 (Figs. 1B and 8A). Additionally, HetOlV2 was transmitted to 2052 from 2072 at sampling spots 1 (Fig. 8A) and 2. Based on RAMS fingerprinting (Supplemental Fig. S8), the genotype of 2052 remained unaltered in all the above cases, which excluded the possibility of the viruses’ presence due to intermixture of recipient and donor hyphae. The transmission of virus(es) was detected in nine other samples, but RAMS analysis indicated that these were contaminated with hyphae of the donor strain, which could be explained by the higher chance of intermingling and overgrowth of hyphae on the 3-month time horizon. Overall, the horizontal transmission rates of the investigated ssRNA viruses between conspecific *Heterobasidion* isolates were 0–33%, which is rather low compared with previously observed rates for dsRNA viruses. Partitiviruses were successfully transmitted in 74% of all tested donor-recipient pairings of *H. parviporum* (Ihrmark et al. 2002; Vainio et al. 2013; Hantula et al. 2020); the transmission frequency of individual viruses ranged 0–100%. The transmission rate of the curvulavirus HetRV6 between *H. parviporum* isolates was 43% (Hantula et al. 2020). The alphapartitivirus HetPV13-an1 was transmitted from its natural host *H. annosum* to 69% of recipient *H. annosum* strains, but the two mitoviruses present in the donor were not transmitted at all (Vainio et al. 2018). The transmission of viruses related to those hosted by our donor isolates has not been studied in *Heterobasidion in vitro*, although there is indirect evidence that ambi-like viruses are being laterally transmitted between *H. parviporum* strains under natural conditions (Sutela et al. 2021).

### 3.10. Effects of virus infections on host growth

Results of the growth rate comparisons are presented in Fig. 8B and C. The naturally virus-free isolate 2052 grew significantly faster than the original virus-hosting isolates 1987, 1989 and 2072 both 3 and 5 dpi (all *p*<0.001). These differences are noteworthy but not necessarily indicative of any mycoviral effects. In fact, 2052 maintained its entire advantage when compared with the cured strains 1987–1W and 2072–1W but performed equally well as 1989–1W (Fig. 8B), which points to the role of inherent phenotypic differences between the tested isolates. In line with the above, no significant differences were found in the growth rates of 1987 or 2072 and their isogenic virus-free counterparts, but 1989–1W advanced significantly faster than 1989 at both points in time (*p*<0.001). The presence of HetTlV1 caused a 19% growth reduction in its host 5 dpi. This reduction indicates that this putative tobamovirus may be affecting essential processes within the host.

No significant differences were observed between 1987 or 1987–1W and the monohyphal cultures. Among the single hyphal cultures, 1987–C1 grew significantly slower than 1987– A2, –A4 and –A5 3 dpi (*p*<0.05), however differences were non-significant 5 dpi. In three comparisons, it was possible to consider the effect of a single virus. Comparing 1987–C1 with –A2, we conclude that HetNlV4 may have a minor positive effect (12%) on host growth, which nonetheless diminished over time (Fig. 8). Confronting 1987–A2 with –A4 and –A5, we suppose that HetNlV2 and HetFV1 are cryptic. Latency is more common among narnaviruses than fusariviruses. Some members of *Fusariviridae* have been associated with hypovirulence in plant pathogenic fungi (Chu et al. 2002; Hao et al. 2018).

The newly infected strain 2052–2072D–1–3M performed significantly worse than its isogenic counterpart 3 dpi (*p*<0.05) and 5 dpi (*p*<0.001). Thus, the introduced HetOlV2 reduced the growth of its new host strain by 20%, despite not affecting its original host strain. The differential response of the two host strains to HetOlV2 indicates that the effects of the virus may be influenced by host-specific factors, such as genetic background, physiological differences or differences in RNAi activity. In such cases, the virus could replicate more efficiently or evade host defenses in the new host strain, leading to reduced growth or other adverse effects (Segers et al. 2007; Hammond et al. 2008; Nuss 2011; Vainio et al. 2015a; Chiba et al. 2016). As stated by Hyder et al. (2013), summarizing data on dsRNA viruses of *Heterobasidion*, “a single mycovirus strain may confer different effects on different host strains”. Members of *Magoulivirus* are considered to establish latent infections with the exception of Fusarium oxysporum ourmia-like virus 1, which exhibited hypovirulence in inoculation experiments on bitter gourd plants (Zhao et al. 2020).

The other isogenic isolates of 2052, infected by various combinations of ambi- and narna-like viruses, also showed some degree of underperformance. 2052–1987D–1 and 2052–1987D–1– 3M were negatively affected both 3 and 5 dpi (all *p*<0.001), with a 5-dpi growth decrease of 13% and 22%, respectively. 2052–1987D–2 suffered a 9% growth reduction 5 dpi (*p*<0.05). Among these three derivatives of 2052, –1987D–1–3M grew significantly slower compared with the other two both 3 and 5 dpi (all *p*<0.05). These data imply that HetAlV19 and HetNlV4 may be somewhat detrimental in coinfection, and their effect is modified by further viruses present in the mycelium. The phenotypic effects of ambiviruses have been investigated in one study (Linnakoski et al. 2021), where the ambi- and ourmia-like viruses of *Armillaria* spp. did not have a major effect on the laboratory growth rate of their hosts either as single or coinfections. Sclerotinia sclerotiorum narnavirus 5 was shown to induce hypovirulence in its host strain harboring five other (+)ssRNA viruses (Hai et al. 2022), which accords with the present findings. As a final point, it should be emphasized that the taxonomic position of a mycovirus is not indicative of its effect on the host (García-Pedrajas et al. 2019).

Taken together, even though some differences were statistically highly significant, the extent of fungal growth reduction was modest across all the comparisons. For reference, HetPV13-an1 reduced the growth of different *H. annosum* and *H. parviporum* isolates by 82–96% under similar experimental conditions (Vainio et al. 2018; Kashif et al. 2019; Hantula et al. 2020). HetPV15-pa1 caused 80–88% growth decrease in *H. annosum* (Kashif et al. 2019). Other partitiviruses and HetRV6 were cryptic or had modest or non-significant effects, either positive or negative to *Heterobasidion* (Vainio et al. 2010; Vainio et al. 2012; Hyder et al. 2013; Kashif et al. 2019; Hantula et al. 2020). Kashif et al. (2019) and Hantula et al. (2020) have shown that coinfecting dsRNA viruses of *Heterobasidion* affect each other’s phenotypic effects on their hosts in an unpredictable manner, i.e. the viral effects are not additive. The lack of any consistent trend for the isolates with multiple infections allows us to speculate that the same holds true for other taxonomic groups of *Heterobasidion* viruses. Detailed studies investigating the effects of each discovered virus in single and coinfections on multiple hosts would be the essential next steps for evaluating their suitability for potential virocontrol applications.

### 3.11. Effects of virus infections on the host proteome profile

The proteomic analyses, utilizing *Heterobasidion annosum* v2.0 (Olson et al. 2012) proteome annotations, resulted in the identification and quantification of 2357 and 1660 protein families, respectively (Supplemental Table S4). The experiment failed to identify any viral proteins. A low-confidence search focusing only on expected viral proteins yielded few potential matches, but these were also found in samples without viruses and were likely false positives. That suggests that either the virus produces very few proteins or the fungus actively suppresses viral transcription/translation.

Although no viral proteins were found in the proteome analysis, a comparison of isolates containing viruses (1987, 1989, 2072) with a virus-free isolate (2052) revealed 510 proteins with statistically significant differences in abundance (differentially abundant proteins, DAPs; *p*<0.05). These DAPs constituted over 40% of the estimated protein levels in isolate 2052. Additionally, the corresponding principal component analysis (PCA) showed a clear distinction between the isolates, with the first principal component (representing 43.4% of the variation) appearing to separate samples based on their viral load (Fig. 9A). The pw comparisons showed relatively low overlap in identified DAPs (*t*-test, *p*<0.05; relative fold change > 1.4), and only 78 proteins showed a similar response to virus presence (Fig. 9B). The subset of proteins that demonstrated an increase in abundance included three enzymes involved in detoxification and stress response (heme peroxidase, DyP-type peroxidase and 2-oxoglutarate/Fe(II)-dependent dioxygenase), proteins participating in targeted protein degradation (DUF431-domain-containing protein and two ubiquitin-conjugating enzymes), various carbohydrate-active enzymes (CAZymes), and an RNA-binding protein (RRM domain-containing protein). In all three virus-infected isolates, an enzyme laccase showed a significant increase in abundance. This finding aligns with prior observations of viral influence on fungal laccase production (Nuskern et al. 2021). Proteins that possibly participated in the fungal response to viruses and showed a significant decrease in abundance included (among others) nine isoforms of glycoside hydrolases, which are believed to play a role in biotic interactions (Bradley et al. 2022), two protein kinases, PKS_ER domain-containing proteins that are involved in secondary metabolism, cytochrome P450 monooxygenases, an enzyme in chitin biosynthesis glutamine-fructose-6-phosphate aminotransferase, proteases, and several proteins with an unknown function.

**Fig. 9.**
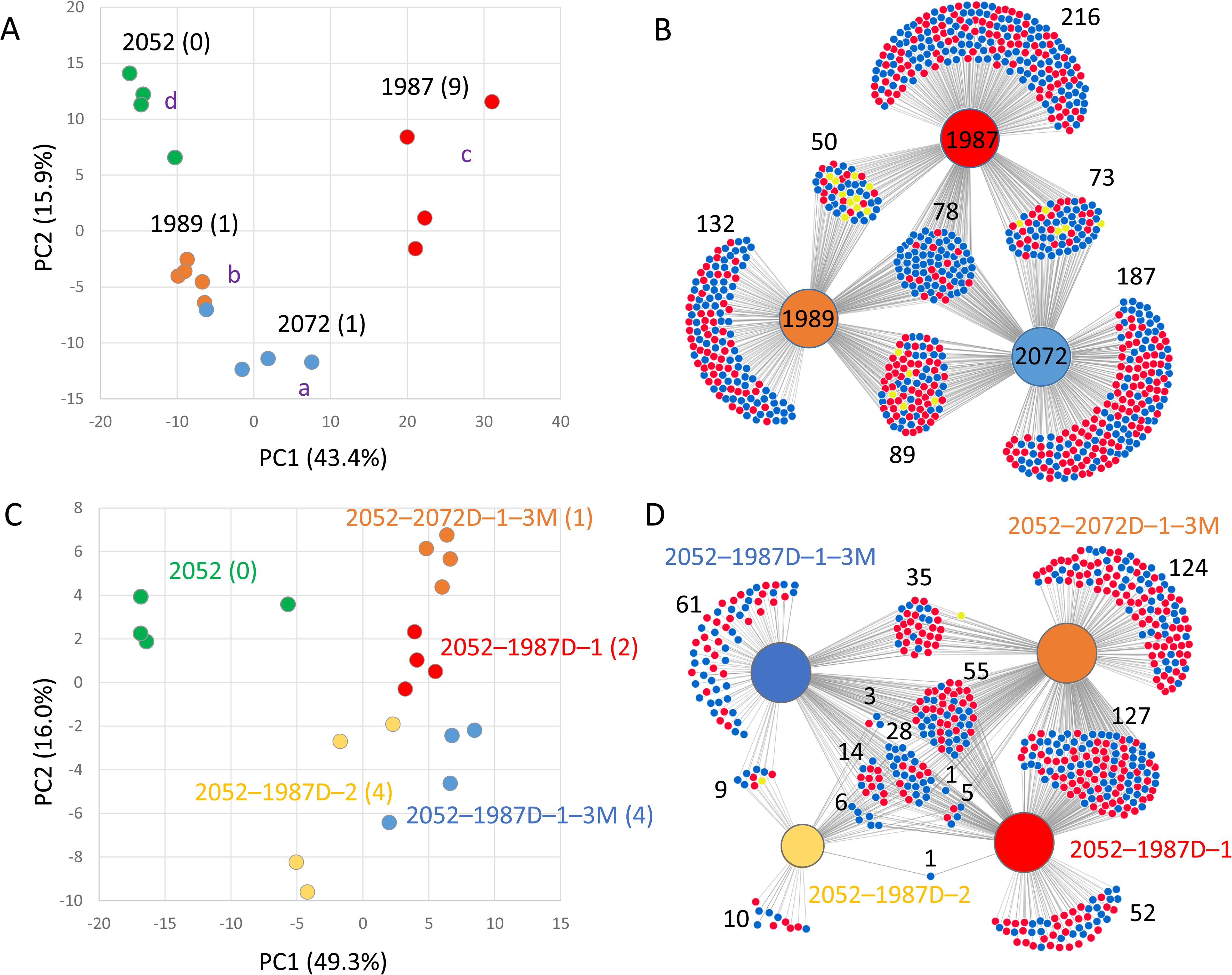
*Heterobasidion* proteome profile reveals distinct signatures in virus-free and infected mycelia. **A** Principal component analysis (PCA) based on 510 differentially abundant proteins (DAPs) identified in four isolates with varying viral content (ANOVA, *p*<0.05, Tukey’s HSD). Different letters indicate statistically significant differences between groups (Conover’s test, *p*<0.05). Numbers in brackets represent the number of different viruses detected in each isolate. **B** Pairwise comparisons of individual isolates to the virus-free isolate 2052, visualized using a DiVenn diagram (Sun et al. 2019). Proteins with a statistically significant change in abundance compared to isolate 2052 (Student’s *t*-test, *p*<0.05; fold change > 1.4) are represented by dots. Red dots indicate proteins with increased abundance, and blue dots indicate proteins with decreased abundance in the compared isolate relative to 2052. Yellow dots represent proteins that showed contrasting responses (increased in one comparison, decreased in another) between isolate pairs. **C-D** Impact of viral infection on the proteome profile: PCA based on all 139 identified DAPs (ANOVA, *p*<0.05, Tukey’s HSD), and the results of pairwise comparisons visualized using a DiVenn diagram (*p*<0.05; fold change > 1.4). All results are based on four biological replicates. For detailed information on protein identifications and statistical analyses, see Supplemental Table S4

In the second set of comparisons, the impact of virus introduction was observed in isolate 2052. Newly infected isolates displayed only 139 DAPs based on ANOVA and Tukey’s HSD test. However, the corresponding PCA successfully grouped the infected isolates as expected (Fig. 9C). Further analysis using consecutive pw comparisons revealed 97 DAPs (*t*-test, *p*<0.05; relative fold change > 1.4; Fig. 9D) with a similar profile in at least three different isolates. Among the DAPs consistently accumulated in all infected isolates were candidates of interest, including a putative ortholog of Prd-6, an ATP-dependent RNA helicase involved in nonsense-mediated mRNA decay, and two proteins involved in ribosome biogenesis: an ATP-dependent RNA helicase involved in the biogenesis of 40S and 60S ribosomal subunits and a ribosomal 60S subunit protein L30. Ribosomal proteins are now a well-established part of fungi–virus interaction (Li 2019), and nonsense-mediated mRNA decay has been found to restrict virus replication in both plants and animals (Popp et al. 2020). DAPs with higher abundance in at least three infected isolates included two cytochrome P450 monooxygenases, an F-box domain-containing protein (involved in targeted protein degradation), and an RRM domain-containing protein (RNA binding). It should be noted that among the 97 filtered DAPs, 72 were also present in the first dataset. Notably, the response to viral presence was similar for 60 of these shared DAPs.

Finally, the proteome of cured isolates and their corresponding infected genotypes were compared. Despite the relatively large sets of DAPs, only 12 DAPs were found in more than one comparison (Supplemental Table S4), and only seven of these showed a similar trend.

Two dioxygenases (Isopenicillin N synthase-like and Fe2OG dioxygenase domain-containing protein) showed significantly higher abundance in cured isolates. Same was found for a nuclear cap-binding protein subunit and a putative autophagy-associated protein (ATG16). Conversely, a lipid metabolism enzyme (3-oxoacyl reductase), an adhesin domain-containing protein, and pyruvate decarboxylase exhibited increased abundance in virus-containing isolates (Supplemental Table S4). The comparison of partially cured isolates 1987 also showed very high diversity with only 14 DAPs showing similar trend in more than two partially cured isolates (Supplemental Table S4). Besides already listed proteins, this dataset identified the putative role of two additional proteins: a Hydantoinase_B domain-containing protein (showing a significant increase in abundance in cured isolates) and a cytosolic Ca2+-dependent cysteine protease (more abundant in infected isolates) that could participate in viral protein degradation.

Taken together, this analysis identified over 500 protein candidates encompassing diverse metabolic and signaling pathways as potential viral response proteins. Approximately 150 of these candidates exhibited the expected trends in abundance changes across at least two of the analyses (Supplemental Table S4).

## 4. Conclusions

Our findings indicate that the virome of *Heterobasidion* populations in Czechia is highly diverse and seems to differ from that in the boreal region. Notably, the most frequently reported virus of the pathogen, HetRV6 was not found among 85 *Heterobasidion* strains from Czechia or Slovakia. Earlier, HetRV6 has been detected in the neighboring countries Austria and Poland (Vainio et al. 2012). Likewise, the *Partitiviridae* family, with 21 members described to date from *Heterobasidion*, was not represented in our datasets although present in the neighboring Poland. However, two previously known and 23 novel ssRNA mycoviruses were discovered. We report the first putative negative-strand RNA virus in *Heterobasidion*. RNA-Seq of a single isolate had clear benefits over RNA-Seq of pooled libraries, such as the discovery of new ambi-like viruses and a narnaviral segment, the completed genome of a virus causing low titer infection and resolved ambiguous bases in certain viral contigs. Some of the tested viruses exhibited a negative impact on the mycelial growth rate and induced considerable alterations in the host proteome profile. Considering both the growth rate test and proteomics, the candidates deemed most suitable for further studies are HetTlV1 and HetOlV2. More extensive studies are needed to evaluate their biocontrol potential.

## Acknowledgements

Computational resources were supplied by the project “e-Infrastruktura CZ” (e-INFRA CZ LM2018140) supported by the Ministry of Education, Youth and Sports of the Czech Republic. We would like to thank Milica Raco (Phytophthora Research Centre, Mendel University in Brno) for guidance in RNA extractions and bioinformatics. We are thankful to Ondřej Hejna (Department of Genetics and Agrobiotechnology, University of South Bohemia in České Budějovice) for support in bioinformatic analysis. Petr Sedlák, Michal Tomšovský, Libor Jankovský and Miloň Dvořák (all Department of Forest Protection and Wildlife Management, Mendel University in Brno) are gratefully acknowledged for invaluable assistance during the collection of *Heterobasidion* isolates.

## Author contribution

All authors contributed to the study conception and design. LBD, MČ and LB conducted experiments. LBD, MČ, MdlP and LB analyzed data. LBD, MČ and LB wrote the manuscript and all authors commented on previous versions of the manuscript. All authors read and approved the manuscript.

## Funding

This research was funded by the Specific University Research Fund of the Faculty of Forestry and Wood Technology, Mendel University in Brno, grant number LDF_VP_2019034. Additionally, it was supported by the European Regional Development Fund, Project “Phytophthora Research Centre”, Reg. No. CZ.02.1.01/0.0/0.0/15_003/0000453 and by Generalitat Valenciana through PROMETEO program, Project CIPROM/2022/21.

## Data availability

The RNA-Seq data have been deposited into the NCBI Sequence Read Archive database with the accession numbers SRR18240564, SRR18240565, SRR18240566 and SRR18961509 under the BioProject PRJNA809936. The mycoviral genomic sequences are available from the NCBI GenBank. The mass spectrometry proteomics data have been deposited to the ProteomeXchange Consortium via the PRIDE (Perez-Riverol et al. 2022) partner repository with the dataset identifier PXD045991.

## Declarations

### Conflict of interest

The authors have no relevant financial or non-financial interests to disclose.

## Supplementary materials

**Table S1.**
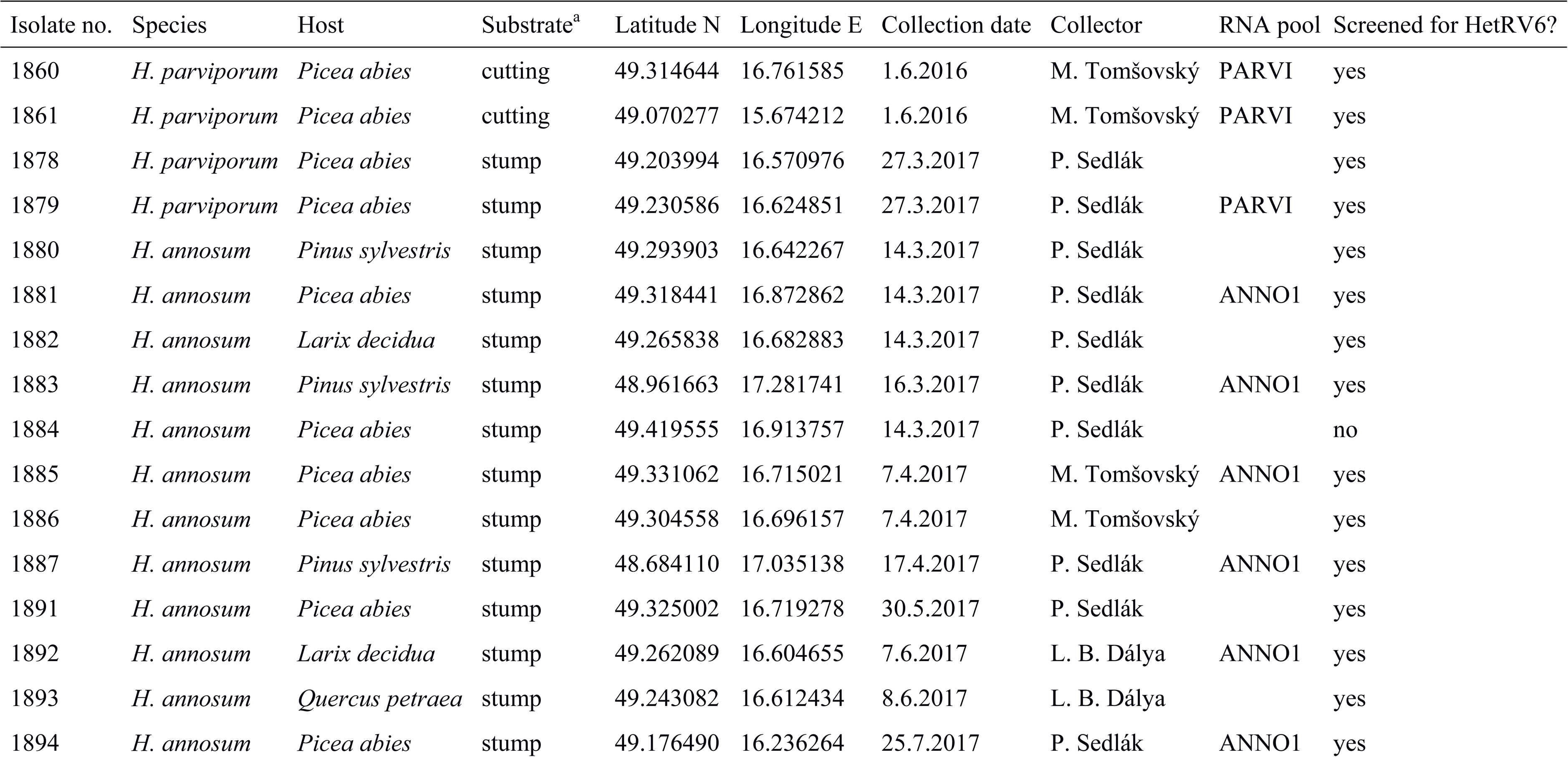
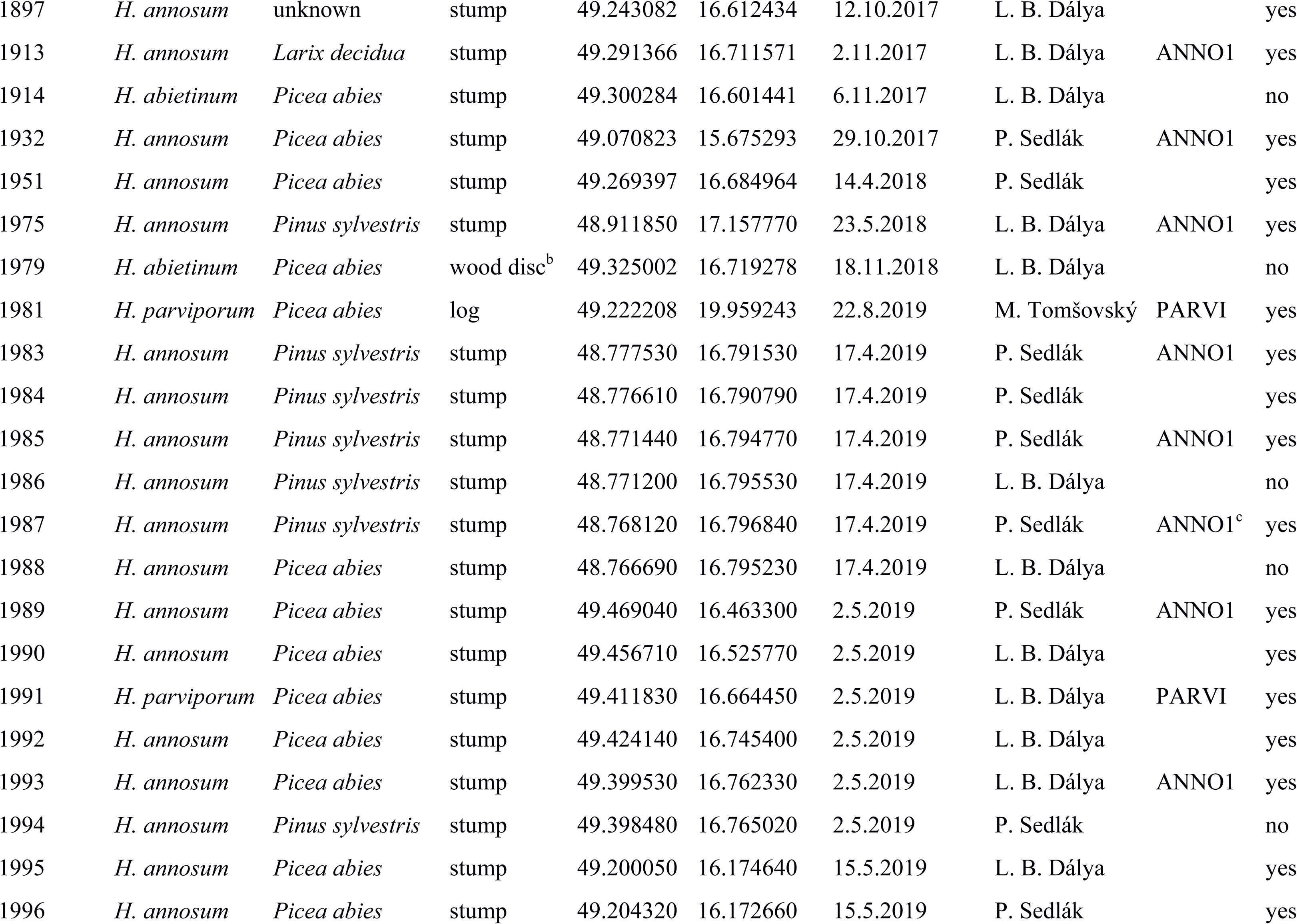
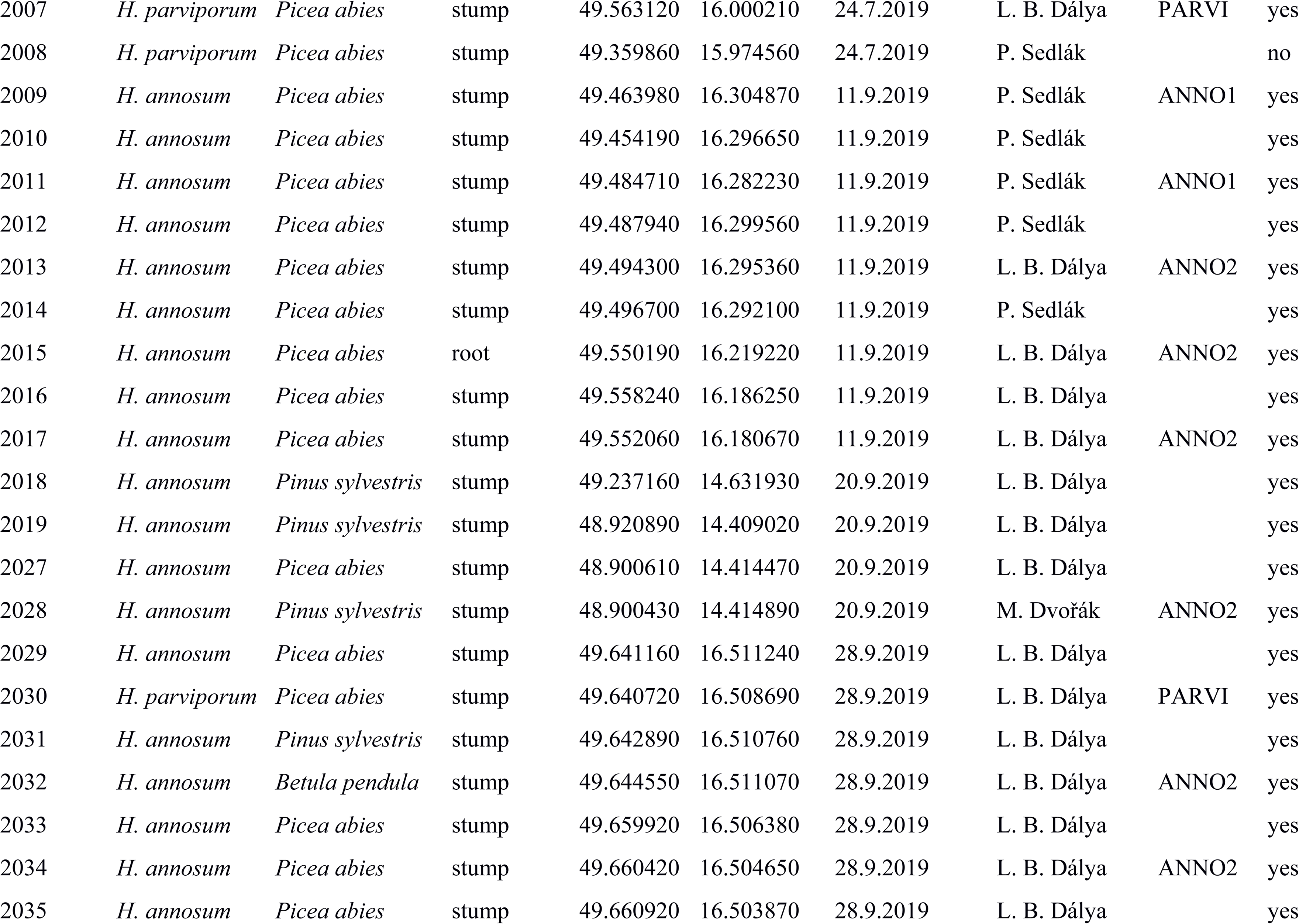
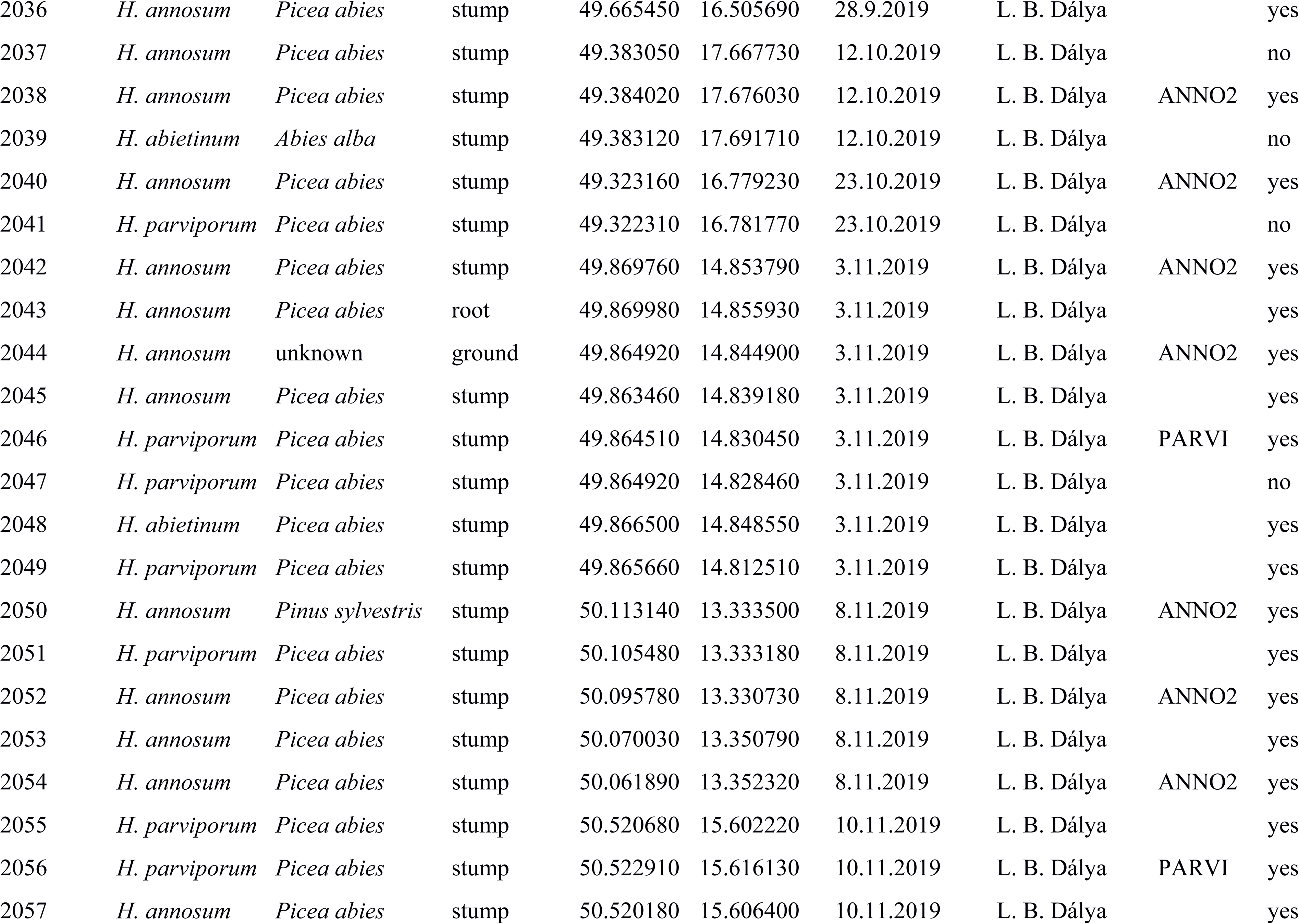
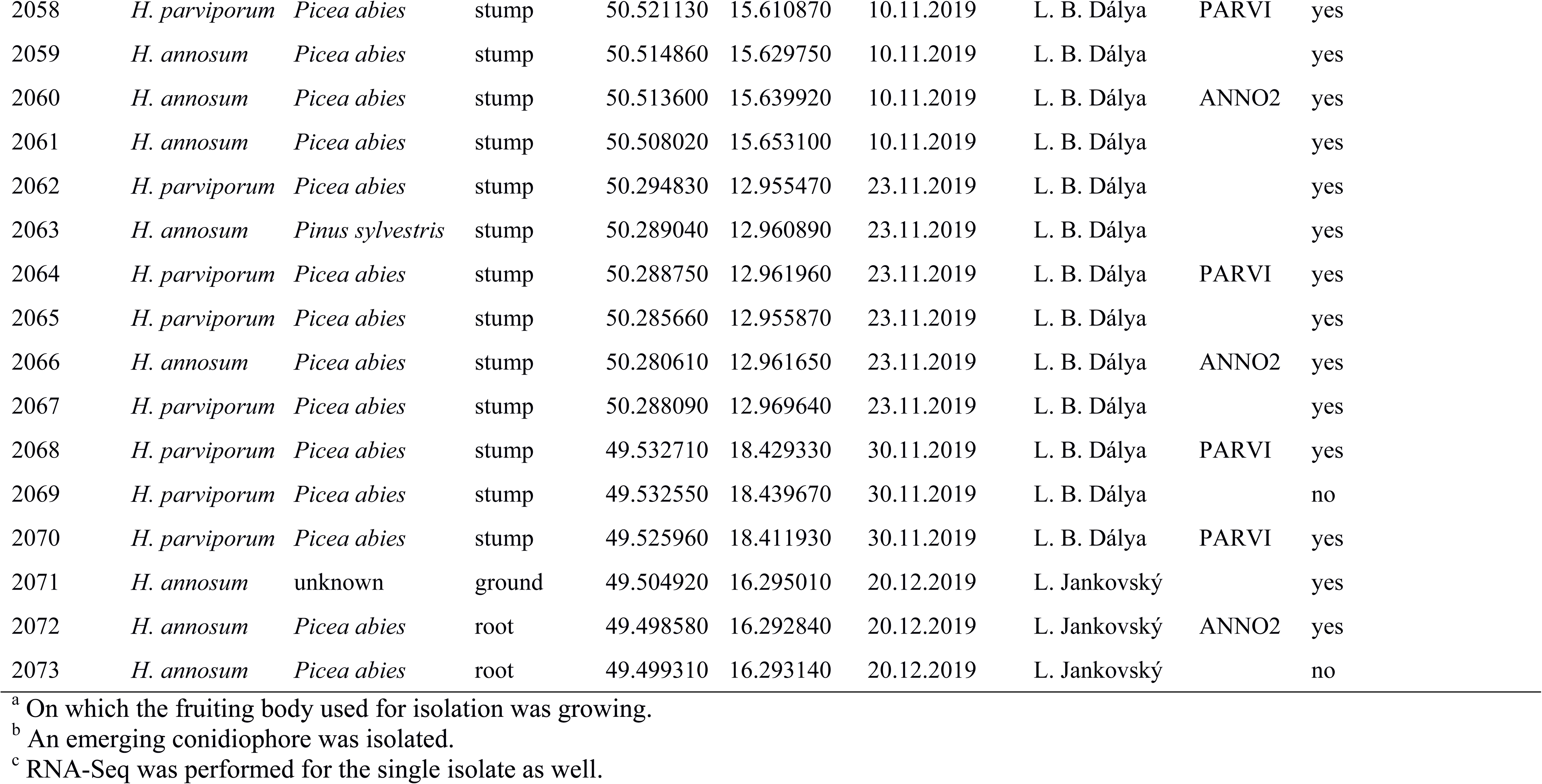
*Heterobasidion* isolates studied.

**Table S2.**
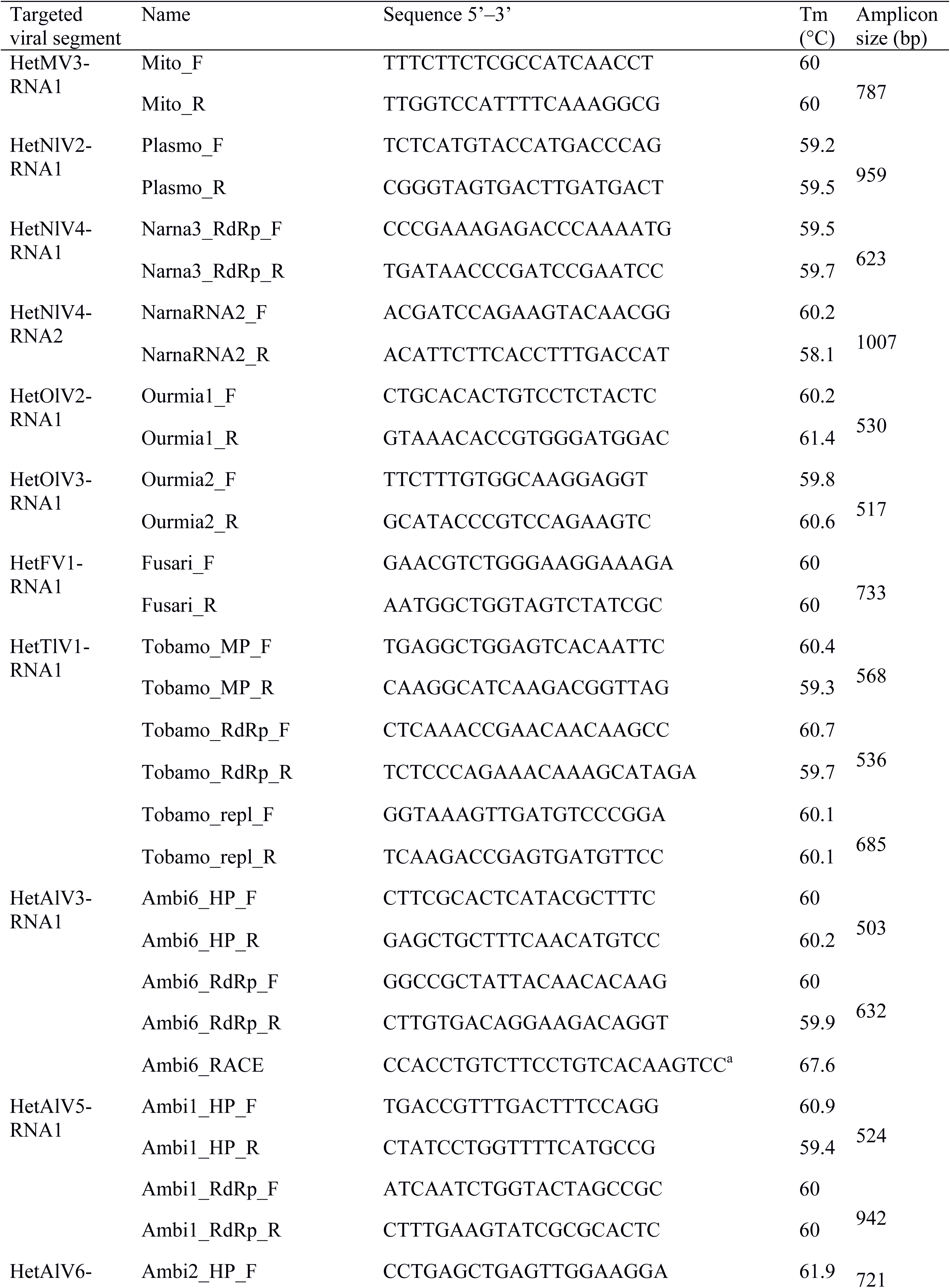
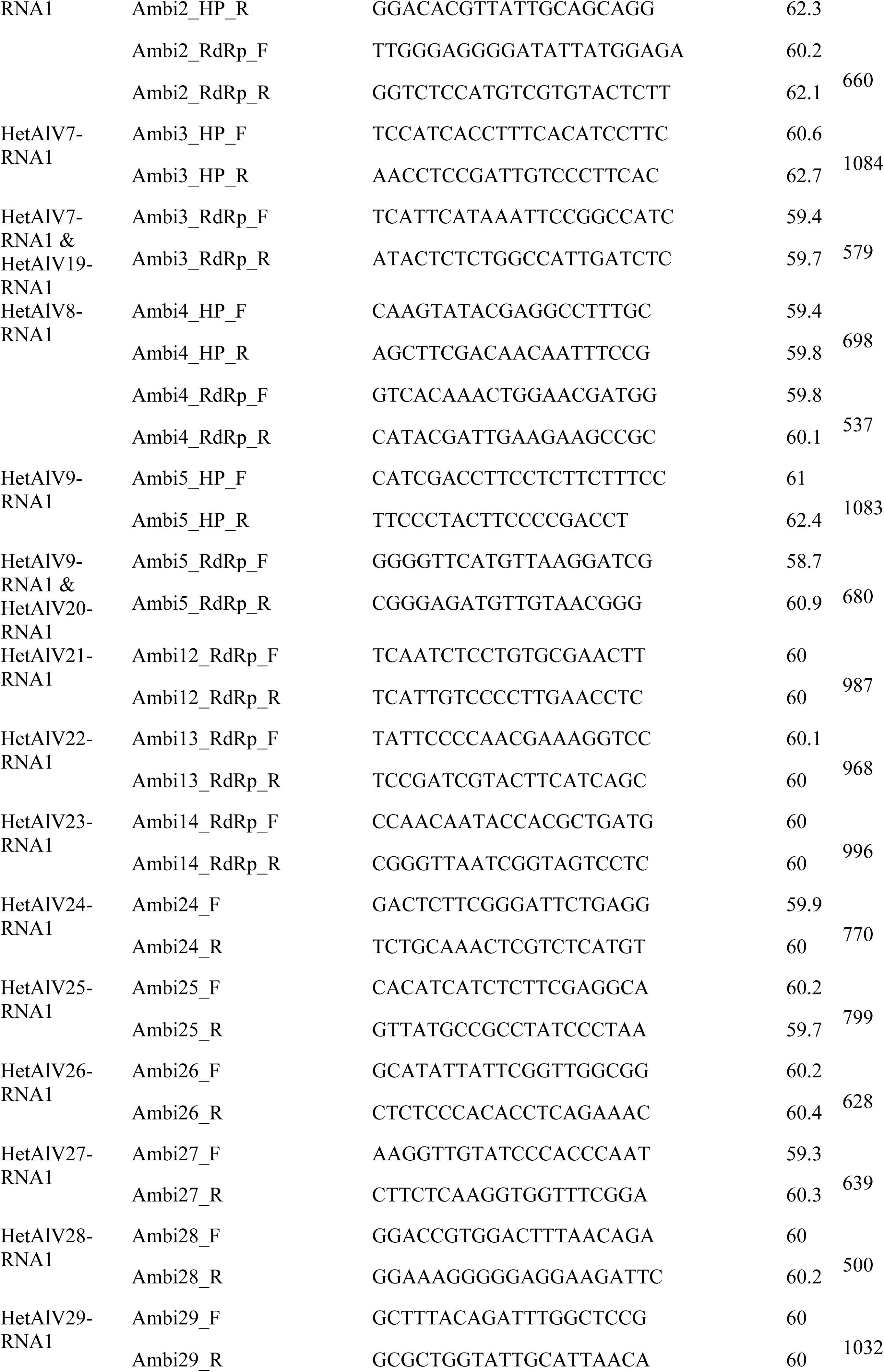
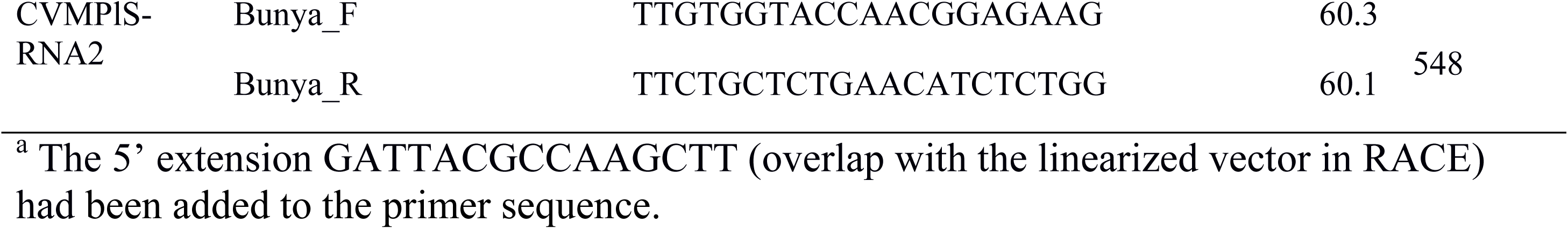
Primers designed and used for the current study.

**Table S3**

Types and genomic position of the ribozymes detected in *Heterobasidion* ambiviruses

**Table S4**

Results of proteomic analyses

**Fig. S1** Confirmation of putative viruses discovered in the RNA-Seq datasets PARVI, ANNO1 and ANNO2. M, molecular size marker (GeneRuler™ 1 kb Plus DNA Ladder, 75– 20,000 bp, Thermo Fisher Scientific, Waltham, MA, USA); 1–2, RT-PCR products amplified by primers Ambi1_HP and Ambi1_RdRp from isolate 2050; 3–8, Ambi2_HP and Ambi2_RdRp from 2028, 2038 and 2050; 9–12, Ambi3_HP and Ambi3_RdRp from 2028 and 2040; 13–16, Ambi4_HP and Ambi4_RdRp from 1983 and 1993; 17–18, Ambi5_HP and Ambi5_RdRp from 2028; 19–20, Ambi6_HP and Ambi6_RdRp from 1991; 21–22, Bunya and Fusari from 1987; 23, Mito from 2009; 24, Ourmia1 from 2072; 25–26, Ourmia2 from 2028 and 2038; 27–29, Plasmo from 1983, 1987 and 1993; 30–32, Tobamo_MP, Tobamo_RdRp and Tobamo_repl from 1989. * The targeted band of 698 bp was purified from the gel prior to Sanger sequencing

**Fig. S2** Pairwise identities of RT-PCR amplicons of mycovirus strains. Values below the diagonal line are nt identities and values above are aa identities

**Fig. S3** Conserved aa sequence motifs I–VI within the RdRp of HetFV1 and selected fusariviruses. LeFV1, Lentinula edodes fusarivirus 1 (QOX06044); PgFV1, Phlebiopsis gigantea fusarivirus 1 (UJT31800); RsFV2, Rhizoctonia solani fusarivirus 2 (QDW92691); SrFV1, Sclerotium rolfsii fusarivirus 1 (AZF86096). Similarity percentages in the MUSCLE alignment were calculated based on Blosum62 score matrix with a threshold of 1

**Fig. S4** Conserved aa sequence motif DEXH within the helicase domain of HetFV1, selected fusariviruses and more distantly related members of *Flaviviridae*. HCV, Hepatitis C virus (2ZJO); JEV, Japanese encephalitis virus (2Z83); DENV2, Dengue virus 2 (2BHR); KOKV, Kokobera virus (2V6I); ZIKV, Zika virus (5GJC); ATP bs, adenosine triphosphate binding site. Similarity percentages in the MUSCLE alignment were calculated based on Blosum62 score matrix with a threshold of 1

**Fig. S5** Conserved aa sequence motifs I–III within the methyltransferase domain of HetTlV1 and selected members of *Virgaviridae*. AbMV1, Armillaria borealis mycovirgavirus 1 (QUD20345); LeTLV1, Lentinula edodes tobamo-like virus 1 (QOX06054); PpTLV, Podosphaera prunicola tobamo-like virus (ATS94406); PvLaT-LV1, Plasmopara viticola lesion associated tobamo-like virus 1 (QIP68002); BSMV, Barley stripe mosaic virus (AAA46336). Similarity percentages in the MUSCLE alignment were calculated based on Blosum62 score matrix with a threshold of 1. The numbers of deleted bases are indicated in gray boxes

**Fig. S6** Conserved aa sequence motifs I–VI within the helicase domain of HetTlV1 and selected members of *Virgaviridae*. Similarity percentages in the MUSCLE alignment were calculated based on Blosum62 score matrix with a threshold of 1. The numbers of deleted bases are indicated in gray boxes

**Fig. S7** Conserved aa sequence motifs I–VIII within the RdRp of HetTlV1 and selected members of *Virgaviridae*. PpTLV, Podosphaera prunicola tobamo-like virus (ATS94407); BSMV, Barley stripe mosaic virus (AAA66600). Similarity percentages in the MUSCLE alignment were calculated based on Blosum62 score matrix with a threshold of 1. The numbers of deleted bases are indicated in gray boxes

**Fig. S8** Banding patterns of random amplified microsatellite (RAMS) fingerprinting with CCA primer. bp, molecular size marker (Thermo Scientific O’GeneRuler Express DNA Ladder, Thermo Fisher Scientific); −, negative control

## Notes

### Competing Interest Statement

The authors have declared no competing interest.

